# Strain-Specific Epistasis Shapes Fitness Landscapes of APOBEC3G Antagonism By HIV-1 Vif Proteins

**DOI:** 10.1101/2025.10.20.683452

**Authors:** Caroline A. Langley, Michelle Lilly, Harmit S. Malik, Michael Emerman

## Abstract

Host immune factors shape viral evolution. The HIV-1 Vif protein counteracts host APOBEC3G (A3G) to ensure productive infection. Using deep mutational scanning (DMS) across two divergent HIV-1 Vif proteins, we systematically mapped and compared the mutational landscapes governing Vif antagonism of A3G. These high-resolution fitness maps reveal core principles of host–virus coevolution. Most missense mutations were strongly deleterious, reflecting pervasive purifying selection. Yet several highly conserved Vif residues at binding interfaces with A3G, RNA, and CBFβ exhibited unexpected mutational tolerance, indicating structural flexibility at these sites. Comparative analysis revealed that strain-specific epistasis dictates Vif–A3G interactions, including adaptive changes at the Vif–A3G interface. Further analysis uncovered striking temporal epistasis, whereby initially adaptive Vif mutations critical for antagonizing hominoid A3G subsequently exhibited reduced fitness. Thus, structural robustness permits adaptation, while epistasis imposes constraints on Vif’s ability to maintain A3G antagonism while remaining responsive to new evolutionary pressures.

**Teaser:** Mapping how HIV-1 evolves to outsmart a key human antiviral protein reveals both unexpected flexibility and evolutionary constraints in viral adaptation.

## Introduction

The interplay between viral proteins and host restriction factors drives a dynamic evolutionary arms race, imposing intense selective pressures on both viruses and their hosts (*1, 2*). Host species evolve mechanisms to recognize and neutralize viral components, whereas viruses evolve countermeasures to subvert these defenses. This reciprocal adaptation fosters continuous cycles of molecular innovation that enhance viral persistence and transmission, while simultaneously shaping host antiviral responses over evolutionary timescales. Such host–virus conflicts have profoundly influenced the evolutionary trajectories of lentiviruses, including Human Immunodeficiency Virus type 1 (HIV-1) and its Simian Immunodeficiency Virus (SIV) precursors in nonhuman primate species.

Among the most potent innate defenses against HIV-1 are those encoded by members of the Apolipoprotein B mRNA Editing Enzyme Catalytic Polypeptide-like 3 (APOBEC3, or A3) family of cytidine deaminases. In primates, this gene family comprises seven enzymes (A3A through A3H), with A3G being the most effective inhibitor of HIV-1 (*3, 4*). The antiviral activity of A3G depends on its incorporation into budding virions and subsequent delivery to target cells, where it localizes to the site of viral reverse transcription. Upon encountering single-stranded viral DNA, A3G deaminates cytidine residues at the second position of 5′-CC dinucleotides (*5, 6*), resulting in G-to-A hypermutation in nascent viral DNA during reverse transcription (*5–8*). These mutations often introduce missense or nonsense changes into the viral genome, severely impairing replication and blocking productive infection. To counteract A3G and other A3 family members, HIV-1 encodes the viral infectivity factor (Vif), a multifunctional protein that prevents A3G incorporation into virions by promoting A3G degradation within the producer cell, thereby neutralizing A3-mediated antiviral activity (*9, 10*). This function is mediated through Vif’s recruitment of several host cofactors: Elongin B (EloB), Elongin C (EloC), and core-binding factor β (CBFβ). Together, these host factors form an E3 ubiquitin ligase complex (referred to as the VCBC complex) that, together with Cullin 5 (Cul5) and Ring-box protein 2 (Rbx2), ubiquitinate A3G and target it for proteasomal degradation, thereby reducing the risk of deleterious hypermutation in the viral genome (*9–15*).

Molecular differences in A3 proteins between hominids and Old-World monkeys have posed significant barriers to the cross-species transmission of lentiviruses, including SIVs and HIV-1 (*16–19*). These interspecies differences have driven the evolutionary diversification of Vif, selecting for mutations that enable antagonism of the distinct A3 proteins found in different primate hosts (*20–22*). Comparative analyses of primate A3G proteins and lentiviral Vif sequences have pinpointed key interaction surfaces and adaptive substitutions at the Vif–A3G interface (*21–23*). One notable example of such adaptation is the emergence of a histidine residue at position 83 in Vif, which arose during the transmission of a recombinant SIV to chimpanzees. This Y83H substitution was critical for enabling Vif from an Old World monkey lentivirus to antagonize hominid A3G following transmission to chimpanzees, thereby overcoming a critical species barrier and facilitating the emergence of SIV_cpz_ and, ultimately, HIV-1 in humans (*22*).

Structural and evolutionary analyses have significantly advanced our understanding of Vif–A3G antagonism. Recent high-resolution cryo-EM structures of HIV-1 Vif bound to human A3G have corroborated and expanded upon comparative evolutionary studies, validating key A3G-interacting residues and uncovering additional contacts, particularly RNA-binding residues that stabilize the Vif–A3G interface through an RNA-mediated scaffolding mechanism (*23–25*). However, even these detailed structural insights provide only a static snapshot of the highly dynamic Vif–A3G coevolutionary process.

Vif sequences from individuals living with HIV-1 exhibit extensive natural variation, including numerous polymorphisms that modulate the antagonism of A3 proteins or other host factors such as PP2A (*26–40*). A recent comprehensive analysis of global HIV-1 *vif* diversity identified sites evolving under either purifying selection or pervasive diversifying selection (*26*), underscoring both ongoing constraints and adaptive pressures acting on this gene. This persistent sequence variability raises critical unanswered questions about Vif’s mutational landscape, its adaptive flexibility, and the impact of natural variation on viral evolution, immune evasion, and pathogenesis. While comparative sequence analyses can provide important insights, they reflect the combined impact of multiple selective pressures acting on Vif, including restriction factors, cofactors, immune responses, genomic constraints (such as overlapping reading frames in the native viral genome), and non-selective founder effects. As a result, evolutionary conservation alone cannot disentangle which constraints are specific to the Vif–A3G arms race. For instance, residues that appear invariant across isolates may be mutationally flexible with respect to A3G antagonism yet conserved because they are required to antagonize other host proteins or overlap with essential viral genes (*e.g.*, *pol*). Conversely, residues that would otherwise be constrained to maintain A3G antagonism may evolve rapidly under competing selective pressures. Thus, although conservation integrates the totality of selective pressures on Vif, it obscures A3G’s specific contribution to these constraints.

Deep mutational scanning (DMS), a powerful high-throughput approach that enables systematic evaluation of the functional consequences of nearly all possible single missense substitutions within a protein (*41–43*), offers a complementary experimental lens. By subjecting large variant libraries to defined selection pressures in pooled assays, DMS directly reveals how individual mutations influence protein stability, intermolecular interactions, and overall fitness. Initially developed to interrogate the functional landscapes of enzymes and essential housekeeping genes (*41–43*), DMS has since been applied to diverse biological systems, offering critical insights into host–virus interactions and mechanisms of viral escape from innate and adaptive immunity (*44–50*). A previous study extended this approach to Vif, using mutagenesis and pooled selection to identify residues required for the degradation of PP2A regulatory subunits, revealing a genetically separable electrostatic interface underlying Vif-mediated G2/M cell-cycle arrest (*32*). These findings highlight that Vif is shaped by multiple, mechanistically distinct selective pressures, including but not limited to A3G antagonism. In this context, applying DMS to Vif provides a means to disentangle the constraints imposed by A3G from the broader set of evolutionary forces captured by sequence conservation.

Here, we applied DMS to systematically investigate *vif* from two distinct HIV-1 Clade B strains in the context of A3G. Specifically, we analyzed *vif* from the HIV-1_LAI_ reference strain, which was used in the cryo-EM structure of the Vif–RNA–A3G complex (*23*), as well as *vif* from the HIV-1_1203_ strain, a primary clinical isolate derived from an individual homozygous for A3H haplotype II, which encodes the most potent antiviral variant of this gene (*51*). Consequently, Vif from the 1203 strain evolved under dual selective pressure to counter both A3G and A3H restriction factors (*31, 51*). We used DMS-derived fitness scores to assess the functional consequences of nearly every missense mutation in both *vif* genes in cells engineered to express only a single APOPEC3 protein, A3G (*3*), thereby isolating A3G-mediated selective pressure. We observed a strong inverse correlation between A3G-mediated G-to-A hypermutation and variant enrichment, confirming that our DMS-derived fitness scores reliably capture Vif’s ability to neutralize A3G. Across both strains, most missense mutations reduced viral fitness, reflecting strong purifying selection. This constraint was particularly pronounced at known interfaces with A3G and CBFβ, consistent with their critical functional roles. Nonetheless, we identified pockets of mutational tolerance, even at residues that are highly conserved across HIV-1 and SIV_cpz_. These include residue 42, which binds RNA, and site 83 within the Vif–A3G “arms-race” interface, where we found mutations that increase the ability of Vif to antagonize A3G in both a lab strain of HIV-1 and in a primary clade A strain, highlighting areas of unexpected structural flexibility in this interaction (*23*). Such unexpected mutational tolerance and adaptive potential in Vif would not have been predicted from structural analyses alone, which provide a detailed architecture of a single snapshot of this highly dynamic arms race, underscoring the complementary insights of both approaches. Comparative analysis of DMS-derived fitness scores for the two Vif genes revealed 70 missense variants at 46 positions with divergent fitness consequences between LAI and 1203 Vif, illustrating how strain-specific selective pressures have shaped the mutational landscape of Vif in distinct ways. Further analysis of this strain-specific epistasis revealed an unexpected finding: the originally adaptive 83H mutation that enabled SIV_cpz_ Vif to counteract hominid A3G now exhibits reduced fitness in some contemporary HIV-1 strains, a striking example of temporal epistasis. Together, these findings offer a comprehensive, high-resolution map of Vif’s evolutionary plasticity, illuminating the trade-offs between maintaining essential antiviral countermeasures and exploring alternative adaptive solutions in the ongoing host–virus arms race.

## Results

### A pooled functional selection assay for HIV-1 *vif* variants

In the HIV-1 genome, the *vif* gene partially overlaps with the *pol* and *vpr* open reading frames. The first 19 codons of *vif* overlap with the 3’ end of *pol*, which encodes the C-terminus of the Integrase (IN) protein (Fig. 1A). To prevent mutations in *vif* from inadvertently disrupting IN function and confounding our readout of A3G antagonism, we removed the *pol*-*vif* overlap (Fig. 1B) as previously described (*16*). We introduced synonymous mutations in the *pol* reading frame that eliminated the endogenous start codon in the *vif* open reading frame (Fig. 1B), as well as three additional in-frame ATG codons. The entire *vif* gene, including a newly introduced start codon, was then repositioned downstream of the *pol* coding region. We also incorporated restriction enzyme sites to enable modular cloning into an otherwise unmodified, replication-competent HIV-1 proviral backbone, as previously described (*16*) (Fig. 1B). To investigate the selective constraints governing HIV-1 Vif antagonism of A3G, we constructed comprehensive DMS libraries encompassing Vif amino acid positions 12-115 (Fig. 1C) (Supplementary data 1 and 2). This region was chosen based on prior studies identifying critical interfaces with A3 proteins (*14, 23, 27, 52–55*). We deliberately excluded the C-terminal domain (residues 116-192), which primarily mediates interactions with the VCBC complex (*12*). This design enabled targeted mutagenesis of functionally important Vif residues that would otherwise be constrained by overlap with *pol*, including residue 15, a well-established determinant of A3G interaction (*23, 56, 57*).

**Figure 1:**
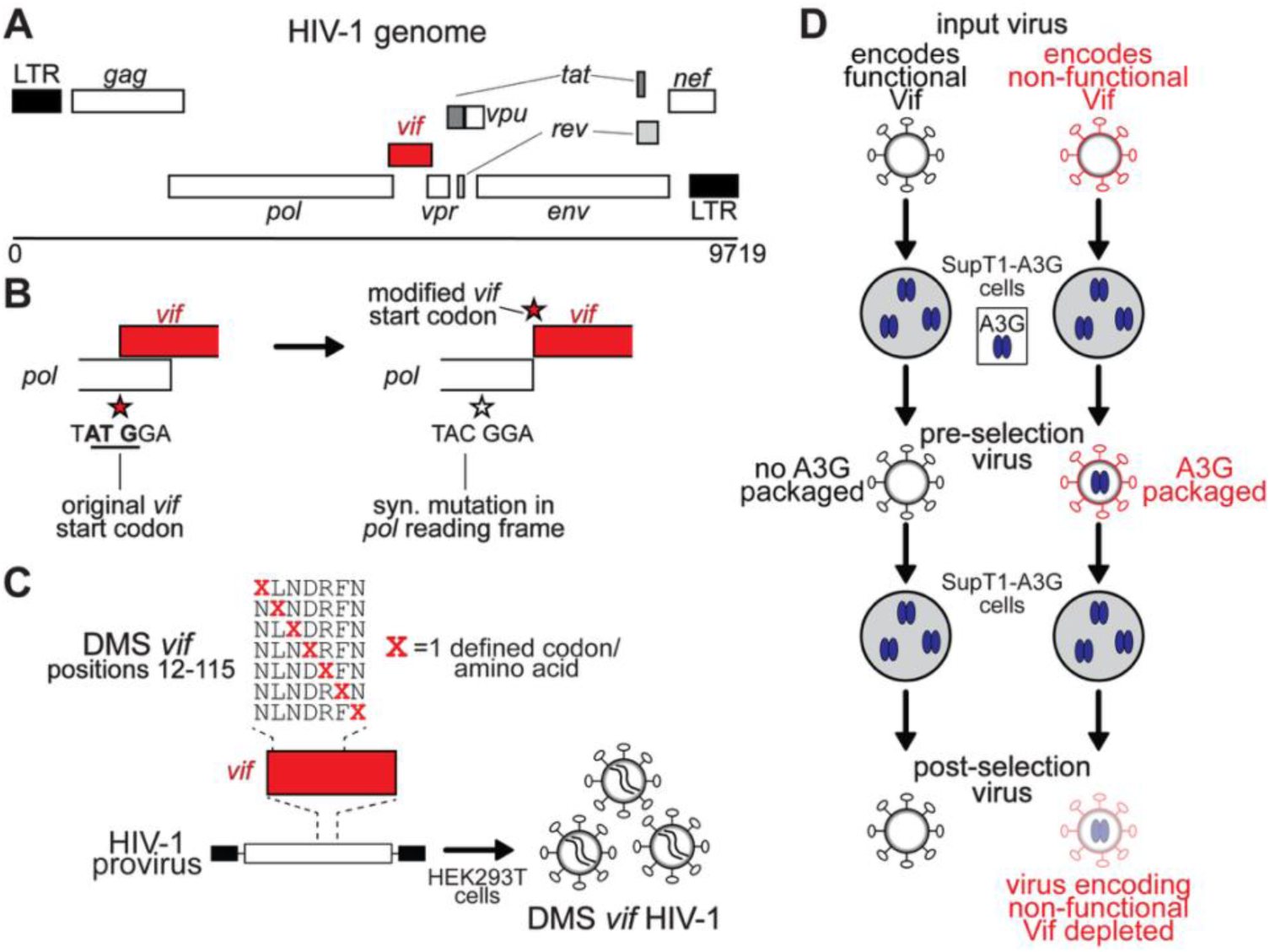
Selection of functional Vif variants through passaging in A3G-expressing cells. **A.** Schematic of the HIV-1 genome with the *vif* gene highlighted in red. **B.** The 5’ end of *vif* overlaps with the Integrase coding region in *pol*. To eliminate this overlap, the endogenous *vif* start codon was disrupted by a synonymous mutation in the *pol* reading frame in a replication-competent HIV-1 proviral backbone. A new *vif* start codon was then inserted downstream of the *pol* reading frame, as described previously (*16, 58*). **C.** Codon-specific DMS libraries were generated across Vif amino acid positions 12-115 from two HIV-1 clade B strains (LAI and 1203). For each amino acid substitution, a single defined codon was used at each site, ensuring uniform representation of missense variants and avoiding over-representation of amino acids encoded by multiple synonymous codons. Codons susceptible to A3G-mediated mutation (*e.g.*, those prone to conversion into stop codons via G-to-A changes) were excluded, thereby minimizing the introduction of nonsense variants. To avoid producing truncated Vif proteins, we also excluded stop codons and Methionine substitutions that could introduce internal translation start sites. The DMS libraries were cloned into the modified proviral backbone and transfected into HEK293T cells to produce infectious DMS virus pools. **D.** DMS virus pools were used to infect SupT1 cells stably expressing physiological levels of A3G (SupT1-A3G) at a low multiplicity of infection (MOI < 0.1). Viruses encoding functional Vif (gray, left) replicate efficiently, whereas viruses encoding non-functional Vif (red, right) incorporate A3G (blue ovals) into virions. Viral supernatant collected 72 hours after infection (“pre-selection”) was used to infect fresh SupT1-A3G cells. During the second passage, virion-packaged A3G deaminates cytidines on minus-strand cDNA, introducing mutations that inhibit replication. Supernatant collected 72 hours later (“post-selection”) was subjected to reverse transcription and deep sequencing to quantify enrichment or depletion of individual Vif variants.

A significant challenge with mutational analyses of Vif is that A3G can induce hypermutations in the viral genome, including within the *vif* gene itself, potentially confounding the interpretation of DMS data. To distinguish DMS-associated substitutions from A3G-mediated mutations, particularly at 5′-CC dinucleotides, which are the preferred deamination target of A3G (*5, 6*), we avoided the conventional error-prone PCR or NNS mutagenesis strategies commonly used in DMS studies. Instead, we devised a codon-specific mutagenesis strategy (Fig. 1C). At each residue of Vif, we introduced alternative amino acids using unique codon substitutions designed to be distinguishable from G-to-A hypermutation events at 5′-CC dinucleotides. Our deliberate missense recoding strategy ensured that all amino acids were equally represented in the variant pool and reduced the likelihood of producing truncated Vif proteins by avoiding ATG codons that could serve as internal translation start sites or stop codons that would lead to premature termination. As a result of our design, fewer than 8% of substitutions in our DMS *vif* libraries generated new 5′-CC dinucleotide contexts that could serve as novel A3G targets; most of these rare instances occurred when introducing tryptophan (TGG) substitutions, where CC dinucleotides cannot be avoided. The resulting DMS *vif* libraries were cloned into the modified, replication-competent HIV-1 proviral backbone and transfected into HEK293T cells to generate pools of infectious HIV-1 virions carrying DMS-mutagenized *vif* variants (Fig. 1C).

To quantify the fitness effects of individual *vif* variants, we developed a pooled, two-passage selection assay in SupT1 cells engineered to express physiological levels of A3G (SupT1-A3G) (*3*) (Fig. 1D). This system provides a genetically homogeneous and reproducible context, minimizes host variability, and enables precise measurement of Vif mutational effects in the presence of a single A3 protein. We used DMS-*vif* HIV-1 virions to infect SupT1-A3G cells at a low multiplicity of infection (MOI) in biological triplicate. Twenty-four hours post-infection, the viral supernatant was replaced with fresh medium to remove viral particles produced before *vif* expression. Seventy-two hours post-infection, we collected the resulting ‘pre-selection’ viral supernatant and used it to infect a fresh batch of SupT1-A3G cells (Fig. 1D). After an additional 72 hours, the ‘post-selection’ supernatant was harvested for deep sequencing (Fig. 1D). In this system, viruses encoding nonfunctional Vif variants that fail to counteract A3G during the initial infection produce progeny virions that incorporate A3G, leading to G-to-A hypermutation and reduced infectivity. Such variants are depleted from the viral population in the post-selection supernatant. In contrast, functional Vif variants that effectively restrict A3G incorporation are enriched in the post-selection supernatant relative to their pre-selection frequencies. Thus, the relative enrichment or depletion of each Vif variant quantifies viral fitness in the presence of A3G. This two-passage selection scheme provides a measure of relative rather than absolute fitness, since even the complete loss of Vif is not expected to result in the complete elimination of those variants from the post-selection virus pool after only one round of selection. Although additional rounds of selection could increase selection stringency, A3G-mediated G-to-A hypermutation would progressively erode the linkage between genotype (Vif variant) and phenotype (viral fitness), which is critical for pooled DMS assays.

To validate the specificity of our assay for selecting functional *vif* variants, we performed two pilot experiments (Fig. 2), each with three biological replicates. First, we compared the replication dynamics of viruses encoding wild-type and a loss-of-function *vif* mutant harboring a premature stop codon at position 40 (Y40*). SupT1-A3G cells were infected with a defined mixture of wild-type (WT) and Y40* viruses. To ensure a measurable dynamic range for enrichment, we used a non-equimolar input ratio (90% Y40*: 10% WT), which allows for detectable increases in WT frequency during selection; in contrast, a 1:1 mixture would compress this dynamic range and limit our ability to quantify enrichment using the selective index. Because A3G-mediated restriction becomes apparent only after two rounds of replication in A3G-expressing cells, we hypothesized that the Y40* variant would be selectively depleted in post-selection, but not pre-selection samples. To quantify selection, we calculated a selective index, defined as the natural logarithm of the ratio of post-selection to pre-selection variant frequencies: *ln(f_post_/f_pre_*), where *f* denotes the frequency of a variant in the indicated population. Frequencies were converted to decimal form prior to transformation. Thus, positive values indicate enrichment, and negative values indicate depletion following selection. For interpretability, we also report the relative percent change, calculated as *(e^(selective^ ^index)^ – 1) × 100*, reflecting the proportional change in representation. Consistent with our hypothesis, the relative abundance of WT and Y40* viruses did not differ significantly between the input and pre-selection (first passage) populations. However, in post-selection samples, the Y40* variant was depleted, corresponding to decreases of 42% and 38% relative to the input and pre-selection populations, respectively (Fig. 2A). These results demonstrate that the assay resolves differences in the relative fitness between functional and non-functional Vif variants. Notably, selection against truncated Vif was not absolute, as viruses harboring the Y40* mutation remained detectable in the post-selection pool, albeit at substantially reduced frequencies. Together, these observations define the assay’s dynamic range and support its suitability for DMS-based analyses of Vif.

**Figure 2:**
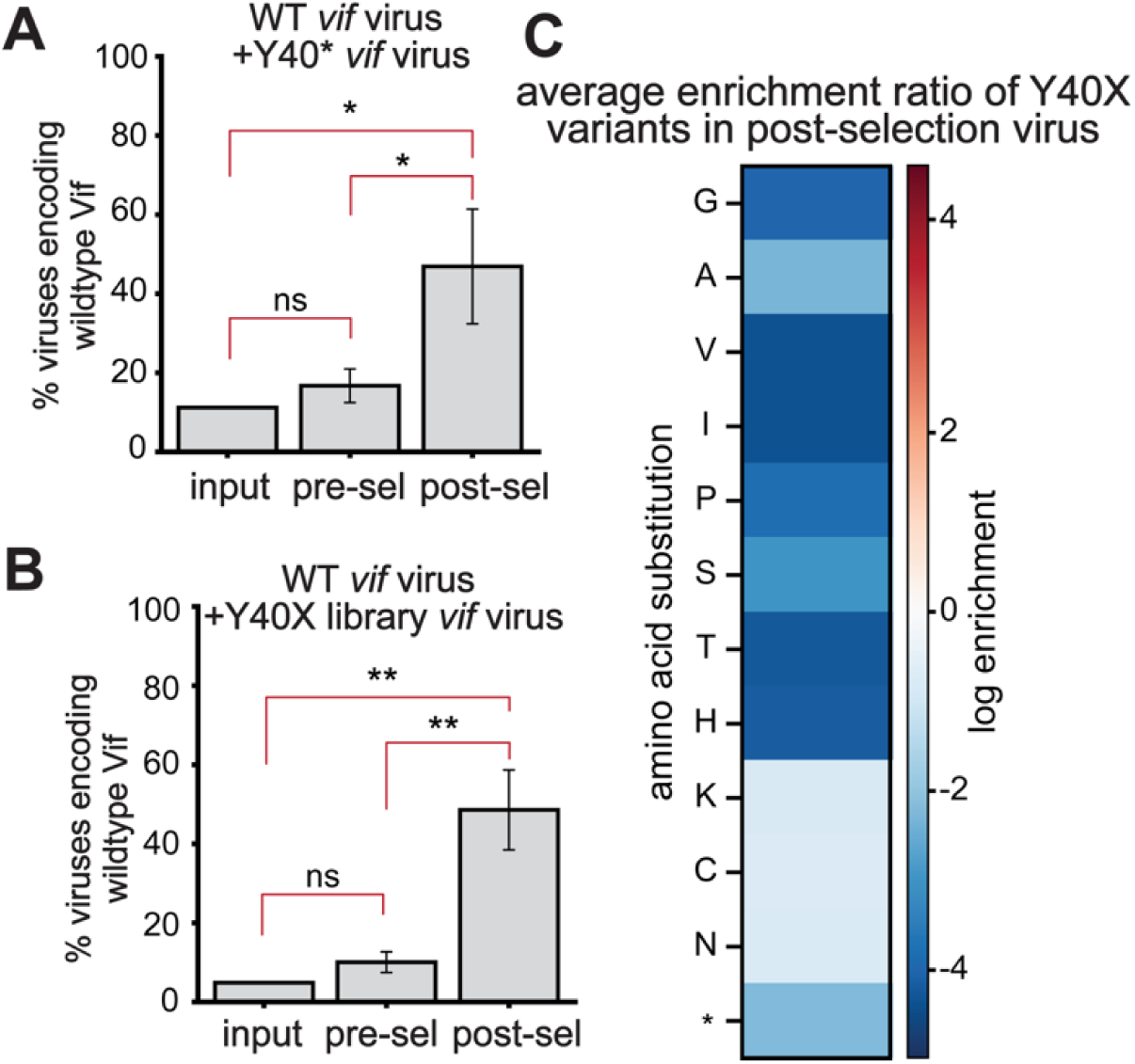
Validation of the selection assay using control virus mixtures. **A.** Viruses encoding wild-type (WT) Vif were mixed with those encoding a premature stop codon mutant at position Y40 (Y40*) of *vif*. **B.** WT viruses were mixed with viruses encoding a representative set of missense variants at position 40 (Y40X) of *vif*. In both cases, the proportion of WT did not change after the first passage (“pre-selection”) but increased only after the second passage (“post-selection”), confirming that A3G-mediated pressure selects for functional Vif variants. Statistical significance was assessed using two-sample t-tests on biological replicates for each pairwise comparison: input vs. pre-selection, input vs. post-selection, and pre-vs. post-selection. Error bars indicate standard deviation across three biological replicates. (* = p < 0.05, ** = p < 0.01, ns = not significant.) **C.** Average enrichment scores for individual missense variants in the Y40X library in the “post-selection” virus pool, averaged across three biological replicates. Blue indicates depletion and red indicates enrichment relative to WT frequency. Amino acids are ordered according to their biochemical properties (*e.g.*, charge, polarity, and hydrophobicity) to facilitate comparison of mutational effects across related residue classes. All missense variants are less preferred than the wild-type Y residue, and several missense substitutions are more deleterious than the Y40* variant.

To further explore the dynamic range of selection, we evaluated a targeted library of missense variants at Vif position 40, a residue known to interact with RNA (*23*), in a second pilot screen. Although this RNA interaction contributes to Vif function, it is not strictly essential, as other Vif residues also contact the same RNA nucleotide (*23*). This experiment enabled us to assess whether our assay could detect more nuanced selective pressures on variants with partial loss of function, rather than the complete loss of Vif function in the Y40* mutant virus. Consistent with expectations, we observed significant depletion of non-wild-type codons at position 40 in post-selection samples, with relative decreases of 46% and 43% compared to the input and pre-selection populations, respectively (Fig. 2B). To visualize the relative fitness of each substitution, we calculated WT-normalized enrichment scores for each variant, defined as *log₂[(f_post_mut / f_pre_mut) / (f_post_WT / f_pre_WT)]* where *mut* and *WT* denote the mutant and wild-type amino acid at the same site, respectively, and plotted these values as a heatmap (Fig. 2C). Notably, several variants exhibited depletion comparable to or greater than that of the Y40* variant, which may reflect dominant disruption of Vif-RNA complex assembly in these variants, resulting in fitness defects that exceed those caused by Vif truncation.

Together, these pilot screens demonstrate that our assay is sensitive to both complete loss-of-function mutations and subtler impairments in Vif activity, thereby validating the robustness of the two-passage selection system. Functional *vif* variants were consistently enriched, whereas those deficient in A3G antagonism were selectively depleted. Importantly, these effects were observed exclusively in post-selection samples, confirming that A3G-mediated selective pressure, rather than experimental artifacts or bottlenecks, drives the observed enrichment patterns.

### Deep mutational scanning of HIV-1 _LAI_ Vif reveals sites of mutational constraint and unexpected tolerance

We then applied the validated two-passage selection assay (Fig. 1) to our complete DMS library, which encompasses ∼2,000 missense variants in the HIV-1 _LAI_ Vif protein. HIV-1 _LAI_ is a well-characterized, lab-adapted strain whose Vif protein was recently resolved in a high-resolution cryo-EM structure bound to A3G, RNA, and the VCBC-Cul5/Rbx2 complex (*23*). To quantify DMS *vif* variant fitness, we calculated log enrichment scores by comparing each variant’s frequency in post-selection replicates (N =3) to its frequency in a pooled pre-selection library combined across replicates to increase coverage and reduce sampling noise (Fig. 3A). This approach allowed the enrichment scores in each replicate to reflect the outcome of an independent selection normalized to a shared baseline. We measured consistency across three biological replicates using Spearman correlation coefficients between post-selection replicates. We observed moderate to high reproducibility (LAI ρ = 0.46–0.66, Supplementary Fig. 1A). These values are consistent with previous gene-wide DMS analyses of HIV-1 *env*, which similarly report moderate to high reproducibility across biological replicates (Pearson R = 0.45-0.76) (*44, 59*). These values indicate consistent selective pressures across replicates but also reflect the inherent noise in large-scale DMS experiments in replicating viral systems. Such variability likely arises from multiple sources, including stochastic infection dynamics at low multiplicity of infection, differences in A3G packaging into virions, and bottlenecks during viral replication and passage. Although these factors introduce noise at the level of individual variants, they are expected to preserve consistent trends across mutation groups and functional regions.

**Figure 3:**
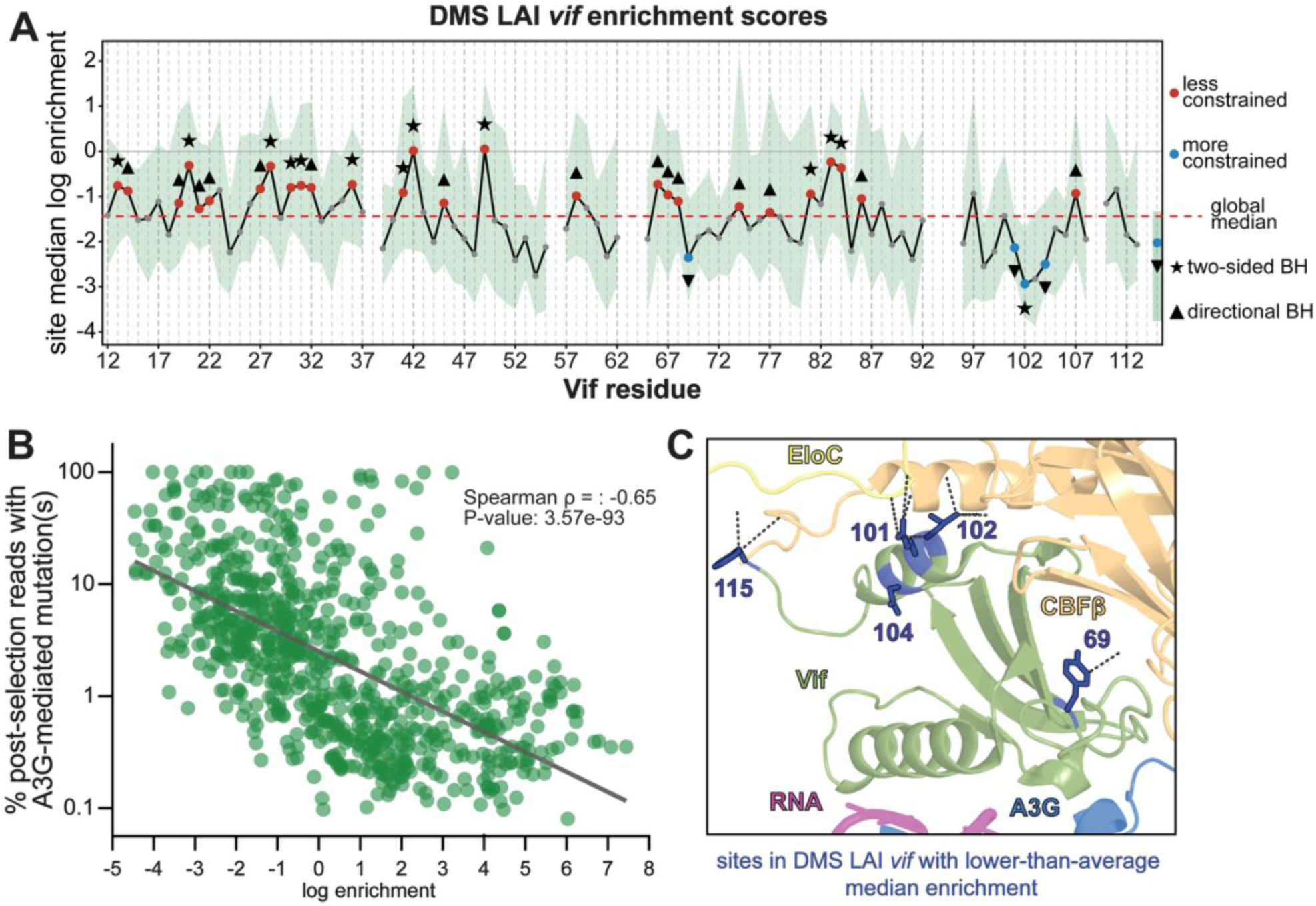
Site-level mutational tolerance of LAI Vif in the presence of A3G. **A.** Plot showing the median log enrichment at each site across all single missense substitutions observed in ≥ 2 post-selection replicates across residues 12–115 of LAI Vif following two rounds of selection in SupT1 cells expressing A3G. The green shading indicates the interquartile range (25%-75%) of variant-level enrichment values at each site. Sites with insufficient variant coverage (<5 missense variants) are not plotted and excluded from statistical analyses. Additionally, sites 38, 56, 63–64, 93–95, 109, and 114 failed library quality control during synthesis and were excluded from analysis. Sites with median enrichment values closer to 0 (solid gray line), or greater than 0, are more tolerant of substitution, whereas more negative values indicate stronger functional constraint. To identify sites whose variant-level enrichment values significantly deviated from the library-wide median (dashed red line), significance was assessed using Wilcoxon signed-rank tests comparing variant enrichment values at each site to the library-wide median (dashed red line). Sites passing a Benjamini–Hochberg false discovery rate threshold (q ≤ 0.10) in either a two-sided test (star) or a directionally consistent one-sided test (triangle) are highlighted in red (less constrained than the global median) or blue (more constrained than the global average). All other sites are shown as gray dots. **B.** Scatter plot comparing the log enrichment ratio of each variant to the percentage of post-selection reads with G-to-A mutations at predicted A3G target motifs. A negative correlation (Spearman ρ = – 0.65, p = 3.57e–93) indicates that higher-fitness variants more efficiently suppress A3G-mediated hypermutation. **C.** Constrained residues (blue dots from panel A) were mapped onto the cryo-EM structure of the Vif–A3G–VCBC–RNA complex (PDB: 8CX0). Dashed lines denote known protein–protein interactions involving these constrained residues.

To enable site-specific interpretation, enrichment scores were further normalized relative to the wild-type amino acid at each position. Wild-type values were set to zero, such that positive scores reflect enrichment, whereas negative scores reflect depletion, relative to the wild-type residue. Median log enrichment scores across the three biological replicates were then computed for all variants at each site (Fig. 3A). This analysis revealed positions in Vif that are broadly tolerant or constrained during replication in the presence of A3G, highlighting regions important for structural integrity, cofactor interactions, and potential sites of adaptive diversification.

To confirm that the DMS-derived enrichment scores accurately reflect the ability of Vif to suppress A3G-mediated mutagenesis, we also carried out an orthogonal analysis. In the absence of effective A3G neutralization, virion-incorporated A3G deaminates cytidine residues at the second position of 5′-CC dinucleotides on the minus-strand cDNA during reverse transcription (*5–8*). A3G’s dinucleotide preference is unique among A3 proteins (*3, 60*), enabling the identification of A3G-mediated G-to-A hypermutation signatures in HIV sequences derived from clinical samples (*61–65*). To leverage this distinctive mutational footprint as an orthogonal measure of Vif functionality, we reanalyzed the same post-selection sequencing data used for enrichment calculations. Because each DMS variant is defined by a fixed set of programmed mutations (Fig. 1C), this approach allowed us to distinguish the encoded variant from additional mutations arising from A3G-mediated deamination within the sequenced region. We mapped known A3G target motifs within the LAI *vif* sequence and quantified the frequency of G-to-A mutations at these positions for each variant. We then compared the proportion of A3G-induced mutations per variant to its log-enrichment score (Fig. 3B). This analysis revealed a strong inverse relationship between A3G-induced mutation burden and variant fitness (Spearman ρ = –0.65). Variants with higher fitness (positive enrichment scores) exhibited reduced frequencies of A3G-induced G-to-A mutations (mean 4.2%), whereas depleted variants (negative enrichment scores) showed significantly elevated frequencies (mean 14.2%). Together, these results independently validate that DMS-derived enrichment scores quantitatively capture Vif’s ability to suppress A3G-mediated mutagenesis.

Notably, we observed substantial variation in enrichment scores among different missense substitutions at the same position, often exceeding the differences in site-level medians (Supplementary Fig. 2). While highly enriched substitutions at several sites may reflect their true adaptive potential, extreme values for individual variants may also arise from assay noise, particularly given incomplete depletion of non-functional Vif variants after a single round of selection (Fig. 2). To mitigate the influence of such stochastic outliers, we therefore focused on site-level median enrichment scores across all missense variants and replicates as a more robust measure of constraint (Fig. 3A; Supplementary Fig. 2). Across the 104 targeted residues in the DMS HIV-1 _LAI_ *vif* library, the overall median log enrichment score was -1.47 (Fig. 3A, dashed red line), indicating that most single missense substitutions reduce viral fitness in the presence of A3G. This pattern underscores the overall mutational intolerance of Vif, consistent with its critical role in counteracting host antiviral defenses.

To identify residues with variant effects that deviate significantly from the global trend (defined as the protein-wide median log enrichment, indicated by the dashed red line in Fig. 3A), we assessed whether the median variant enrichment at each site differed from the overall protein-wide median. Specifically, we used Wilcoxon signed-rank tests to compare the distribution of variant-level enrichment values at each site to the global median across all variants. Because this test evaluates the full distribution of variant effects at each site, statistical significance depends not only on the median enrichment value but also on the consistency (*i.e.*, variance) of effects across substitutions; thus, sites with similar median enrichment values can differ in significance if the spread of variant effects differs. Two complementary versions of the test were applied: a two-sided test to detect significant deviations from the global median, and, independently, a one-sided (directional) test to detect consistent enrichment or depletion. P-values from each test were independently corrected for multiple comparisons using the Benjamini–Hochberg procedure (*66–69*), applying a 10% false discovery rate (FDR) threshold (q ≤ 0.10). In practice, all sites passing the two-sided FDR threshold also passed the directional threshold, indicating that detected deviations were consistently directional; final significance calls additionally required reproducible identification in at least two independent replicates. This criterion integrates statistical significance with reproducibility across biological replicates. Given the directional nature of DMS enrichment scores, sites with consistently negative enrichment values relative to the protein-wide median are interpreted as under stronger-than-average functional constraint, whereas sites with enrichment values significantly higher than the global median are considered relatively more mutationally tolerant, even when their absolute enrichment values remain negative.

Using this criterion, we identified five positions under strong functional constraint in the DMS LAI *vif* library (Fig. 3A, blue dots; Fig. 3C). At these sites, most substitutions were markedly depleted, indicating reduced fitness relative to the wild-type residue. In the recently resolved cryo-EM structure of the Vif–VCBC–RNA–A3G complex (*23*), four of these positions – residues 69, 101, 102, and 115 – directly contact CBFβ (Fig. 3C). Additionally, three of these residues (101, 102, and 104) fall within the previously characterized ^96^T(Q/D/E)x_2_Px_2_ADx_2_(I/L)^107^ motif, which is essential for A3G neutralization(*70*). Residue 69 has also been implicated in A3G interaction (*52, 71*), with a Y69A substitution shown to abolish Vif-mediated antagonism of A3G (*71, 72*). Consistent with these prior observations, our DMS data revealed a strong depletion of the Y69A variant in the LAI *vif* library (Supplementary Data 3). Although Y69 does not directly contact A3G in this cryo-EM structure (*23*), it lies adjacent to W70, a residue critical for stabilizing the Vif–A3G interface (*20–22, 57, 73*). Together, these results indicate that highly constrained residues in Vif are concentrated at structurally and functionally critical interfaces, including CBFβ binding and regions proximal to the A3G interaction surface. Even in the absence of direct contact, residues such as Y69 likely contribute to A3G antagonism by maintaining local structure, positioning neighboring residues, or promoting assembly of the VCBC complex.

We also identified 26 positions in the DMS LAI *vif* library with significantly higher median enrichment scores, indicative of relatively increased mutational tolerance (Fig. 3A, red dots; Fig. 4A). Although most of these sites still had negative median enrichment scores, indicating that most substitutions remain deleterious, their fitness effects were less severe than at more constrained regions of Vif. This pattern indicates that these positions are subject to weaker functional constraints or greater structural and mechanistic flexibility. Despite their apparent mutational tolerance, several of these sites are highly conserved across Group M HIV-1 strains (Supplementary data 4). For example, residue 13 is conserved as valine in 99.6% of sequences, and residue 14 is conserved as aspartate in 99.8% of sequences. Notably, both residues fall within the overlapping reading frame shared between *vif* and the C-terminal tail of *pol*-encoded IN (Fig. 1A) in native HIV-1 genomes. Because this *vif-pol* overlap was removed in the proviral construct used for the DMS library, the observed mutational tolerance at these positions may reflect the intrinsic flexibility of Vif in the absence of overlapping genomic constraints, whereas their conservation in circulating HIV-1 sequences may be driven primarily by constraints on the *pol* reading frame rather than direct selection on Vif function in A3G antagonism.

**Figure 4:**
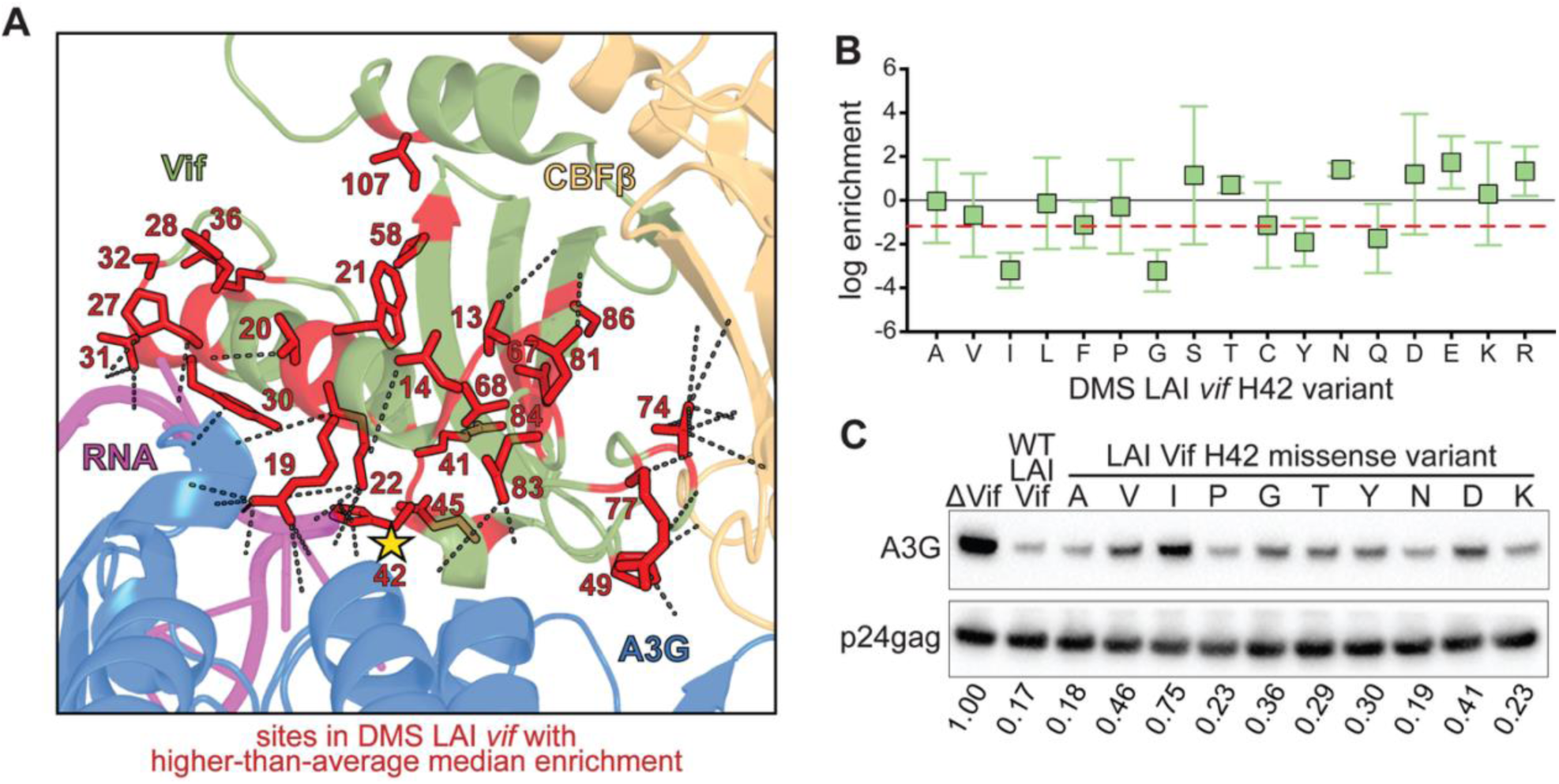
Mutational tolerance at Vif residue H42 reflects both structural flexibility and functional retention in A3G antagonism. **A.** Sites with significantly higher-than-average median log enrichment ratios (from Fig. 3A, red dots) mapped onto the LAI Vif–A3G–VCBC–RNA cryo-EM structure (PDB: 8CX0). Dashed lines indicate known protein–protein and protein–RNA interactions. Residue H42, located at the RNA-binding interface, is highlighted with a yellow star. **B.** Log enrichment ratios from the DMS LAI *vif* library for all single amino acid substitutions at position 42. The black line marks the wild-type histidine (H) enrichment. The library-wide median enrichment is represented by the dashed red line. **C.** Western blot analysis of A3G incorporation into virions following co-transfection of HEK293T cells with FLAG-tagged A3G and individual LAI Vif variants containing single missense substitutions at residue 42. Error bars indicate standard deviation across three biological replicates. Numbers below each blot lane represent the relative intensity of virion-incorporated A3G, normalized to wild-type Vif (H42). Lower levels of virion-incorporated A3G correspond to more effective Vif-mediated restriction.

Fourteen of the 26 mutationally tolerant residues map to established interfaces with A3G, RNA, and/or CBFβ (Fig. 4A, dashed lines) (*23*). The presence of mutational tolerance at these interface residues is unexpected, given their presumed functional importance. One possible explanation is that Vif has evolved structural robustness that permits amino acid variation at these positions without complete loss of function. To investigate this possibility, we focused on position 42, which is perfectly conserved as a histidine (H) residue across HIV-1 and SIV_cpz_ strains. H42 is a critical residue that binds RNA that bridges Vif and A3G (*23*). Despite its conservation and critical importance in Vif interactions, this site appeared highly mutationally tolerant in the DMS LAI *vif* dataset (Fig. 4A, star; Fig. 4B, red line marks the library-wide median). Notably, several substitutions (N, D, and R) exhibited relative enrichment and are accessible via single-nucleotide changes, highlighting the potential for adaptive variation even at canonical interface residues.

To assess whether this tolerance reflects true functional flexibility, we performed an orthogonal A3G packaging assay to assess the functional impact of individual missense variants at HIV-1 _LAI_ Vif position 42 (Fig. 4C). Single missense variants at residue 42 were cloned into a lentiviral expression vector and co-transfected into HEK293T cells together with FLAG-tagged A3G, as described previously (*23*). Seventy-two hours post-transfection, virions were harvested, and A3G incorporation was quantified by western blot (Fig. 4C). Functional outcomes from this virion-incorporation assay largely recapitulated DMS-derived fitness measurements (Fig. 4B). Substitutions with the lowest enrichment scores, such as H42I and H42G (−3.20 and -3.22, respectively; Fig. 4B), led to markedly elevated A3G packaging relative to wild-type Vif (Fig. 4C), indicating substantial loss of A3G-antagonizing function. Moderately depleted variants like H42V and H42Y (average log enrichment values of -0.68 and -1.91, respectively, Fig. 4B) also showed partial functional defects (Fig. 4C). In contrast, H42A, H42P, H42K, and H42N (average log enrichment ratios of -0.04, -0.29, 0.29, and 1.41, respectively) excluded A3G nearly as effectively as wild-type Vif (Fig. 4C). A subset of variants (*e.g.*, H42T and H42D) showed discordant behavior: despite enrichment in the DMS assay, they failed to efficiently exclude A3G in the packaging assay. This discrepancy may reflect partial functional impairments that are not fully resolved in pooled selection, context-dependent effects, or experimental noise. Overall, these results confirm that position 42 tolerates numerous missense substitutions that antagonize A3G, despite its strict evolutionary conservation as a histidine residue. These findings suggest that conservation at this site may reflect historical contingency or overlapping functional constraints, rather than a strict biochemical requirement for A3G antagonism.

### Vif constraints and adaptive potential differ between two divergent HIV-1 strains

To investigate how evolutionary pressures have shaped the mutational landscape of Vif, we compared DMS profiles from LAI with those of a divergent HIV-1 clade B strain, 1203. Although their *vif* genes share 169 of 192 residues (88% identity; Fig. 5A), the strains differ markedly in origin and antiviral activity. HIV-1_LAI_ is a laboratory-adapted strain that efficiently antagonizes A3G but exhibits limited activity against A3H (*58*). In contrast, HIV-1_1203_ encodes a Vif protein that antagonizes both A3G and A3H (*58*), reflecting adaptation to a host environment with potent A3H activity. Consistent with this, residues 45-63 and 90-93 have been identified as key determinants of A3H antagonism in 1203 Vif (*74, 75*).

**Figure 5:**
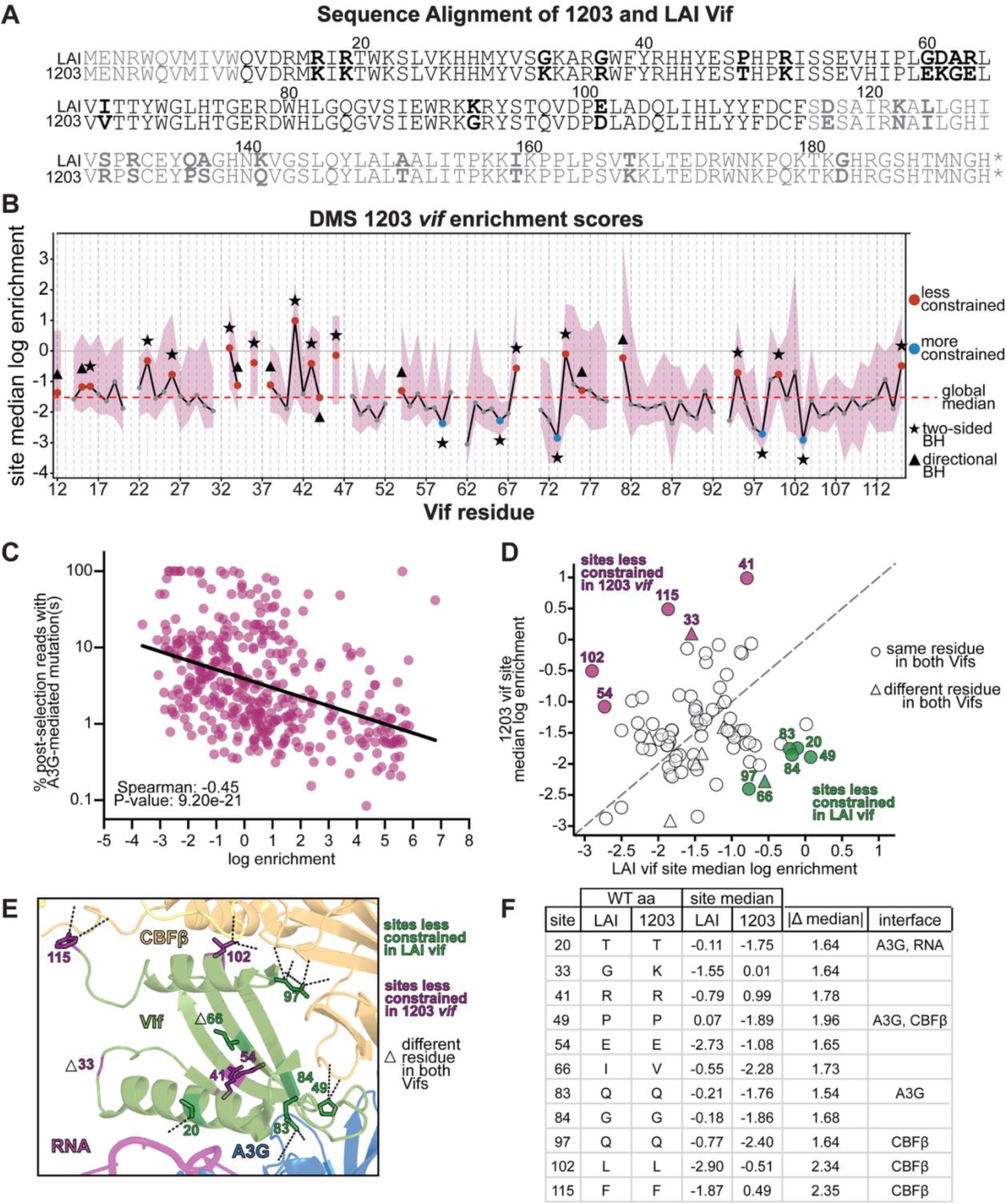
Comparative DMS reveals strain-specific constraints and adaptive divergence between LAI and 1203 Vif proteins. **A.** Amino acid sequence alignment of HIV-1 LAI and 1203 Vif proteins. Divergent residues are bolded in black. Positions not included in the DMS libraries are shown in gray. **B.** Plot showing the median log enrichment at each site across all single missense substitutions observed in ≥ 2 post-selection replicates across residues 12–115 of 1203 Vif following two rounds of selection in SupT1 cells expressing A3G. The purple shading indicates the interquartile range of variant-level enrichment values at each site. Sites with insufficient variant coverage (<5 missense variants) are not plotted and excluded from statistical analyses. Additionally, sites 21, 35, 45, 47, 53, and 93 failed library quality control during synthesis and were excluded from analysis. Sites with median enrichment values closer to 0 (solid gray line), or greater than 0, are more tolerant of substitution, whereas more negative values indicate stronger functional constraint. To identify sites whose variant-level enrichment values significantly deviated from the library-wide median (dashed red line), significance was assessed using Wilcoxon signed-rank tests comparing variant enrichment values at each site to the library-wide median (dashed red line). Sites passing a Benjamini–Hochberg false discovery rate threshold (q ≤ 0.10) in either a two-sided test (star) or a directionally consistent one-sided test (triangle) are highlighted in red (less constrained than average) or blue (more constrained than average). All other sites are shown as gray dots. **C.** Scatter plot comparing each variant’s log enrichment ratio with the frequency of A3G-mediated G-to-A mutations at predicted target motifs. A significant negative correlation (Spearman ρ = –0.45, p = 9.30e–21) supports the use of enrichment scores as a proxy for A3G antagonism. **D.** Comparison of site-level median log enrichment scores between LAI and 1203 DMS *vif* datasets. Each point represents a residue, with axes showing the median enrichment in each strain. Sites are colored if they exhibit large between-strain differences in mutational tolerance (|Δ median| ≥ 1.5 log units), with green indicating sites less constrained in LAI and purple indicating sites less constrained in 1203. Circles denote positions with the same wild-type residue in both strains, and triangles denote positions where the wild-type residue differs. The dashed line indicates equal median enrichment between strains. **E.** Structural mapping of sites with large between-strain differences onto the Vif–A3G–VCBC–RNA cryo-EM structure (PDB: 8CX0). Sites are colored as in panel D. Dashed lines indicate known protein–protein and protein–RNA interaction interfaces. **F.** Table summarizing sites with large between-strain differences in median enrichment, including wild-type residues in LAI and 1203, site-level median enrichment values, absolute differences, and associated interaction interfaces.

We performed three replicates of our DMS selection assay using the 1203 *vif* sequence (Fig. 5B; Supplementary Fig. 4A). We found a moderate concordance between different 1203 replicates, with Spearman correlation coefficients (ρ) ranging from 0.43 to 0.51 (Supplementary Fig. 1B), consistent with the HIV-1_LAI_ DMS (Supplementary Fig. 1A) and previously published DMS analyses of HIV-1 genes (*44, 59*). In contrast, concordance between the HIV-1_LAI_ and HIV-1_1203_ DMS datasets was much lower, with Spearman correlation coefficients (ρ) ranging from 0.00 to 0.16 (Supplementary Fig. 1C). These findings reflect genuine differences in the mutational fitness landscapes of Vif between the two strains. Using the same normalization as for HIV-1_LAI_, we calculated enrichment scores based on the median across all three replicates. We then compared the resulting enrichment scores with A3G-specific G-to-A mutation frequencies (Fig. 5C). As with DMS HIV-1_LAI_ *vif* (Fig. 3B), we observed a moderate inverse correlation between A3G-induced mutation burden and variant enrichment (Spearman ρ = –0.45), supporting that enrichment scores in the HIV-1_1203_ dataset similarly reflect Vif-mediated antagonism of A3G.

We identified 5 constrained sites in 1203 Vif (59, 66, 73, 98, 103) (Fig. 5B, blue dots; Supplementary Fig. 4). Although we also previously found 5 constrained sites in the LAI Vif DMS dataset, there is no direct overlap between these sets of residues. Four of five sites that are constrained in LAI Vif (101, 102, 104, 115) are not constrained in 1203 Vif (constraint at residue 69 could not be reliably assessed in the 1203 Vif dataset). Similarly, none of the 5 constrained sites in 1203 Vif were found to be constrained in the LAI Vif dataset (although residues 98 and 103 approached this threshold). Instead of overlapping constraints at the individual residue level, we observe that constraint is conserved at the level of local structural regions, with neighboring residues exhibiting similar patterns (*e.g.*, residue 103 in 1203 Vif versus residues 101,102, and 104 in LAI Vif). Two of the constrained sites in 1203 Vif fall within the CBFβ interface (sites 73 and 98). This is similar to the LAI DMS dataset, in which four of five constrained sites (sites 69,101, 102, and 115) map to this interface.

We also identified 18 mutationally tolerant sites in 1203 Vif (Fig. 5B, red dots; Supplementary Fig. 4). Like the LAI dataset, a subset of tolerant sites in 1203 localize to functional interfaces, including those involved in A3G and RNA binding (6/18 in 1203; 9/26 in LAI) and CBFβ interaction (3/18 in 1203; 5/26 in LAI). Four of the 18 tolerant sites in 1203 Vif are also tolerant in the LAI DMS dataset (residues 36, 41, 68, and 74); all four sites encode the same wild-type amino acid in both LAI and 1203 Vif proteins. Of these, only site 74 lies within a known interface (CBFβ).

We next compared the enrichment profiles of the LAI and 1203 *vif* DMS libraries at the site level. For each residue, we calculated the median log enrichment for each strain and computed the difference in site-level medians between strains (Δmedian). Sites were then categorized based on the direction and magnitude of this difference: “not significantly different”, “less constrained in LAI”, or “less constrained in 1203”. Globally, the two Vif proteins exhibited nearly identical levels of constraint, with median log enrichment scores of –1.47 for LAI and –1.51 for 1203. This similarity indicates that overall selective pressure is comparable between strains despite differences at individual sites. To focus on sites with the most pronounced differences, we restricted our analysis to sites with an absolute between-strain median difference (|Δ median|) ≥ 1.5, corresponding to the upper tail of the observed distribution (75th percentile = 1.11) (Fig. 5D). Nine of these eleven residues correspond to sites where LAI and 1203 encode identical wild-type amino acids (Fig. 5A; 5D–E, circles; 5F), suggesting that these differences in constraint are not driven by primary sequence differences but instead reflect strain-specific optimization of Vif function under distinct selective pressures. This result demonstrates that differences in the Vif fitness landscape can arise independently of primary sequence variation, highlighting the role of evolutionary contingency and strain-specific epistasis in shaping mutational effects across genetic backgrounds.

Four of the 11 sites with significantly divergent mutational tolerance between LAI and 1203 Vif map to residues that directly contact CBFβ (*23*) (Fig. 5E). Given the central role of CBFβ in stabilizing Vif and facilitating A3G antagonism (*12, 76, 77*), this divergence suggests that Vif has fine-tuned its interaction with CBFβ along strain-specific evolutionary trajectories. Of these four sites, one (residue 20) falls within the shared A3G–RNA binding interface, one (residue 49) lies within the A3G-binding interface, and one (residue 83) is a well-characterized residue within the “arms race interface” of Vif–A3G interactions (*20, 22, 57*) (Fig. 5E). The concentration of sites with divergent mutational tolerance at these functional hotspots supports the idea that multiple genotype-dependent solutions can independently preserve A3G antagonism. This divergence may reflect differences in the broader selective environment experienced by each strain. For example, adaptation to A3H in the 1203 background may indirectly reshape constraints on shared interfaces involved in A3G antagonism. Collectively, these results underscore Vif’s evolutionary plasticity and its capacity to accommodate strain-specific adaptations while safeguarding core functions such as A3G antagonism.

Although site-level analyses provide a broad view of mutational constraint, they can mask important differences in the functional impact of individual missense substitutions. To resolve these finer-scale effects and identify candidate sites for further analysis, we compared the mutational tolerance of all missense variants between the LAI and 1203 *vif* DMS libraries. For each variant, we calculated the mean log-enrichment ratio across biological replicates for each strain. Inter-strain differences were then quantified as Δlog enrichment, defined as *log(LAI) – log(1203)*. Thus, positive values indicate variants that are more tolerated in LAI, whereas negative values indicate greater tolerance in 1203. For interpretability, these values were expressed as linear fold changes (*2^(Δlog^ ^enrichment)^)*. Using this framework, we identified 70 variants across 46 unique Vif residues with significantly divergent functional effects between the two strains, defined as values exceeding 1.5 standard deviations from the mean of the Δlog enrichment distribution (Fig. 6A, colored points). We note that this analysis is intended to highlight regions and interfaces enriched for divergent variant behavior, rather than to draw conclusions from individual variant-level measurements. Among these 46 residues, three map to the A3G-binding interface (blue points), one maps to the RNA-binding interface (pink point), six contribute to both A3G and RNA interactions (red points), three contribute to both A3G and CBFβ interactions (yellow points), and 11 interact with CBFβ (orange points) (*23*).

**Figure 6:**
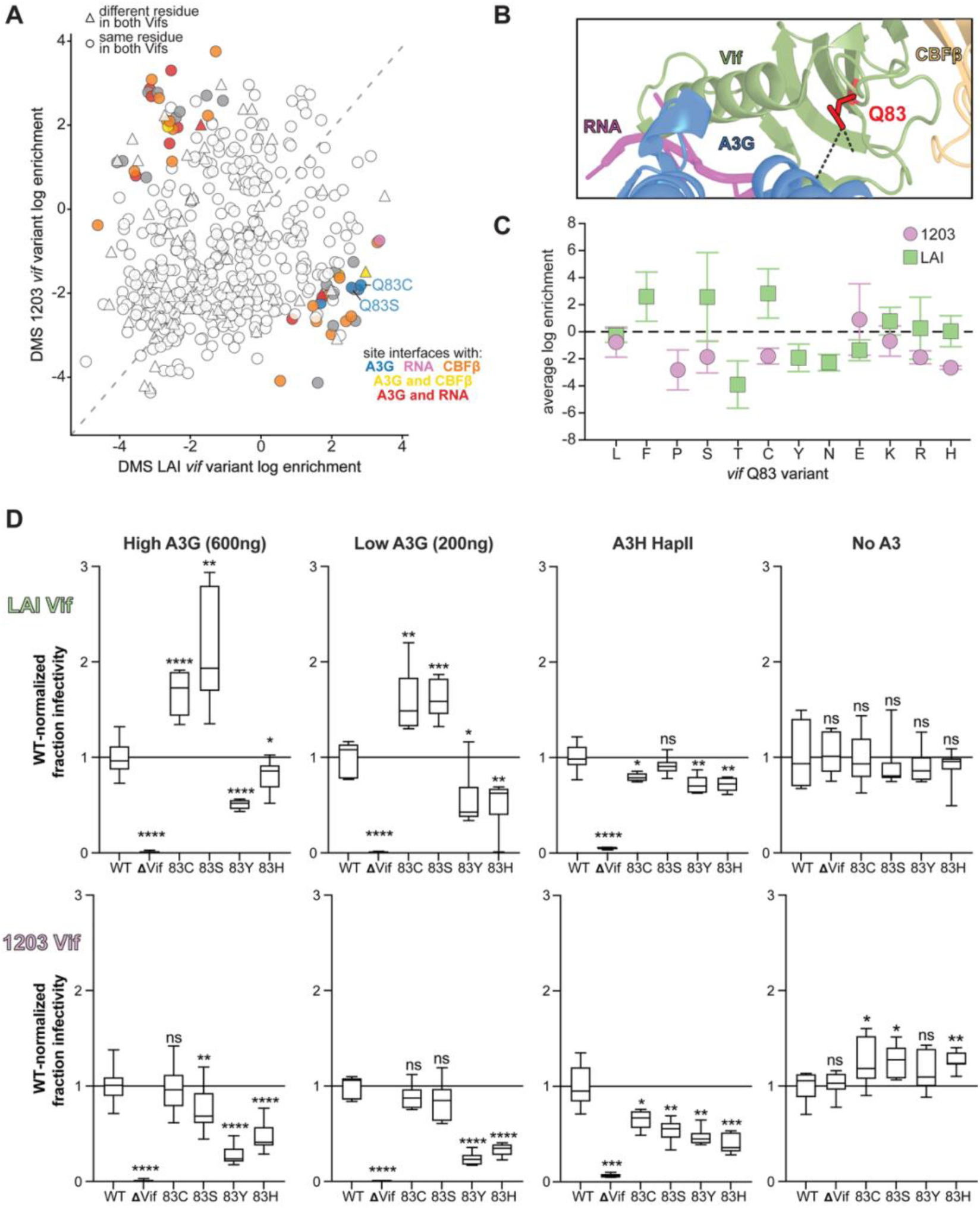
Strain-specific fitness effects of Q83 variants reveal divergent adaptive landscapes in LAI and 1203 Vif. **A**. Scatter plot comparing average log enrichment scores of all single amino acid Vif variants between the LAI and 1203 DMS libraries. Variants with strain-specific fitness effects (≥1.5 standard deviations from the mean of the log[LAI/1203] distribution) are colored according to their mapped interaction interface: A3G (blue), RNA (pink), CBFβ (orange), A3G and CBFβ (yellow), or A3G and RNA (red). Q83C and Q83S are highlighted. **B.** Structural depiction of Q83 (red) in the LAI Vif–A3G–VCBC–RNA cryo-EM complex (PDB: 8CX0), showing its location at the A3G-binding interface. **C.** Average log enrichment scores for all Q83 substitutions in LAI (green squares) and 1203 (purple circles) DMS *vif* libraries. The dashed line at 0 indicates the enrichment score of the wild-type residue. **D.** Infectivity assays for replication-incompetent, VSV-G pseudotyped HIV-1 viruses encoding Q83 variants in the LAI or 1203 Vif background. Infectivity was measured in SupT1 cells expressing high (600 ng of plasmid) or low (200 ng of plasmid) A3G, A3H hapII (200 ng of plasmid), or in the absence of A3 proteins (no A3). Fold changes in infectivity are shown relative to wild-type Vif within each condition, with values for A3G and A3H conditions additionally normalized to the corresponding mean infectivity in the no A3 control. Error bars represent the standard deviation across biological replicates. Statistical significance was assessed using Welch’s t-test, comparing each variant to wild-type Vif: ns = not significant (P > 0.05); *P ≤ 0.05; **P ≤ 0.01; ***P ≤ 0.001; ****P ≤ 0.0001.

Among these sites, we focused on position 83. This site was the only A3G-interacting site with divergent mutational tolerance between LAI and 1203, both at the site level (Fig. 5D-F) and at the variant level, with multiple missense variants exhibiting divergent functional effects between the two *vif* backgrounds. Prior studies have shown that the ancient Y83H substitution was pivotal for cross-species transmission of a precursor SIV into hominids, ultimately enabling the emergence of HIV-1 (*22*). Consistent with its pivotal role in adaptation to hominids, analysis of *vif* sequences from four representatives of each HIV-1 Group M subtype, including circulating recombinant forms, revealed that position 83 most commonly encodes histidine (H) in 68.5% of sequences, followed by glutamine (Q) in 29.9%, and asparagine (N) in 1.4% (Supplementary data 4). In contrast, clade B HIV-1 viruses display a different preference: Q is strongly favored (93.0%), whereas H (5.8%) and N (0.8%) occur only rarely (Fig. 7A; Supplementary data 5). LAI and 1203 are both clade B strains that encode a Q at this site.

**Figure 7:**
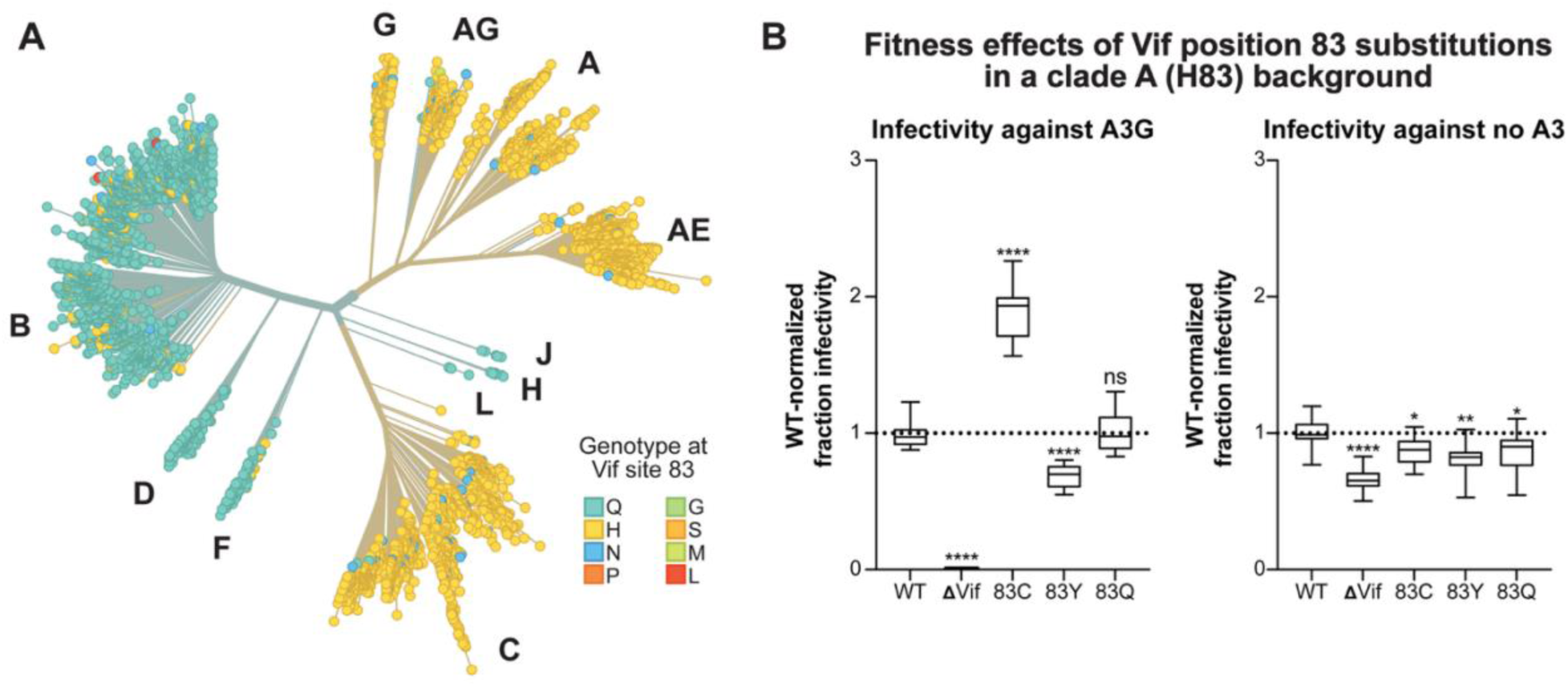
Context-dependent fitness effects of Vif position 83 substitutions in a clade A background. **A.** Phylogenetic tree of HIV-1 Vif sequences colored by the amino acid identity at position 83. Major clades are labeled, and tips are colored according to residue identity. Data were retrieved on October 15, 2025, from Nextstrain’s LANL-HIV-DB view (https://nextstrain.org/groups/LANL-HIV-DB/HIV/genome). **B.** Infectivity of Vif position 83 variants in a clade A Vif background (HIV-1_Q23-17_ strain), in which histidine is the wild-type residue. Viruses encoding wild-type (WT), ΔVif, or the indicated substitutions (H83C, H83Y, H83Q) were tested for infectivity in the presence of A3G (left) or in the absence of A3 proteins (right). Infectivity is normalized to WT, with values additionally normalized to the corresponding mean infectivity in the no A3 control. Statistical significance was assessed using Welch’s t-test, comparing each variant to wild-type Vif: ns = not significant (P > 0.05); *P ≤ 0.05; **P ≤ 0.01; ***P ≤ 0.001; ****P ≤ 0.0001

Consistent with earlier studies, we found that the Q83Y variant, a reversion to the ancestral state prior to adaptation to antagonize hominoid A3G, was strongly depleted in the DMS LAI *vif* dataset (average log enrichment ratio -1.93; Fig. 6C); coverage for this variant was insufficient in the 1203 DMS dataset to make a firm conclusion. Unexpectedly, the historically adaptive Q83H variant (the variant found in SIV_cpz_) exhibited reduced fitness in both strains, with a mild defect in LAI (log enrichment score of - 0.16) and a pronounced defect in 1203 (−2.68) (Fig. 6C). To confirm these effects, we performed independent infectivity assays with pseudotyped viruses encoding representative missense variants in either the HIV-1_LAI_ or HI-1_1203_ Vif background (both inserted into an HIV-1_LAI_ proviral backbone lacking *env*). For each background, variant infectivity was normalized to the corresponding wild-type Vif. We included a ΔVif virus, which does not express a functional *vif* gene, as a negative control. To distinguish A3G-specific effects from broader effects on Vif stability or function, we performed these infectivity assays in the presence of high (600 ng plasmid) or low (200 ng plasmid) A3G, A3H hap II (the most active allele of A3H), or in the absence of any A3 protein (“no A3”) (Fig. 6D).

Our analyses of mutations with divergent fitness effects between LAI and 1203 Vif highlighted two notable substitutions at site 83: Q83C and Q83S. Q83C emerged from the DMS LAI *vif* dataset as the most enriched variant at this position (average log enrichment 2.83). Although Q83C has only been reported in one of 54,228 *vif* sequences in the HIV Sequence Database (Los Alamos National Laboratory, (*78*)), it was also previously shown to enhance Vif antagonism of a divergent A3G in an *in vivo* experimental evolution study (*79*). The second-most enriched variant in the LAI Vif DMS dataset was Q83S (average log enrichment: 2.57). In contrast, both Q83C and Q83S were depleted in the DMS 1203 *vif* dataset, with average log enrichment scores of -1.80 (Q83C) and -1.87 (Q83S) (Fig. 6C). To validate these strain-specific fitness effects, we included these substitutions in our independent infectivity assays with pseudotyped viruses, allowing us to assess the full impact of each substitution across the entire viral lifecycle (Fig. 6D). In the LAI Vif background, both Q83C and Q83S conferred significantly enhanced antagonism of A3G compared to wild-type Vif, as expected from the LAI Vif DMS analyses (Fig. 6D). However, in the presence of A3H hap II, Q83C showed no measurable advantage over wild-type, while Q83S exhibited a modest reduction in infectivity, indicating that the beneficial effects of these substitutions are specific to A3G antagonism. Conversely, in the 1203 Vif background, neither Q83C nor Q83S conferred any advantage over wild-type, while both substitutions were deleterious in the presence of A3H. Together, these results demonstrate that mutations that enhance A3G antagonism in one genetic background can be constrained in another, consistent with epistatic effects shaping Vif function.

As expected, consistent with their depletion in the DMS dataset, both Q83Y and Q83H exhibited significantly reduced A3 antagonism relative to the respective wild-type Vif proteins in both strain backgrounds (Fig. 6D). Importantly, these defects were also observed in A3H antagonism, indicating that the functional impairment due to Q83Y or Q83H is not A3G-specific despite the location of this residue within the A3G-binding interface; instead, this suggests a more global decrease in Vif protein abundance or function. Although it is not surprising that Q83Y leads to loss of Vif function against human A3 proteins, the reduced infectivity of Q83H is especially surprising, given that H83 was initially pivotal for enabling Vif to overcome the restriction imposed by hominoid A3G (*22*). These findings are consistent with temporal epistasis, where subsequent evolutionary changes in Vif from clade B HIV-1 strains have reshaped functional constraints at this site, such that substitutions that were once tolerated or advantageous now impair antagonism of multiple A3 proteins, including A3H.

If this model were correct, we would predict that the Q83H substitution would be less deleterious in non-clade B viruses, where histidine is the predominant residue at this position (Fig. 7A). To test this idea, we introduced both 83Y and 83Q substitutions into a clade A Vif background (HIV-1_Q23-17_ strain, in which H83 is the wild-type residue; Fig. 7B) (*80, 81*). As observed for LAI and 1203 Vif, the 83Y substitution resulted in a significant decrease in infectivity (Fig. 6D). In contrast, H83 and Q83 supported viral fitness to a similar extent in the HIV-1_Q23-17_ strain background, indicating that the Q83H substitution is not inherently deleterious but instead exhibits context-dependent effects. Together, these findings suggest that while the initial Y83-to-H83 transition facilitated adaptation to hominoid A3G, both H83 and Q83 can be functionally equivalent in certain genetic backgrounds, as exemplified by the clade A HIV-1_Q23-17_ virus. However, the functional impairment of Q83H in clade B Vif, coupled with its low frequency in these viruses, points to a lineage-specific constraint that disfavors histidine at this position. These findings highlight the dynamic nature of host–virus coevolution, demonstrating how residues that were once critical for cross-species adaptation can become selectively disfavored due to epistatic interactions with subsequent lineage-specific mutations.

We also found that the H83C substitution conferred a significant fitness gain in the clade A Vif (Fig. 7B), indicating that this effect is not restricted to lab-adapted strains of HIV-1 or to HIV-1 Vif proteins in which Q83 is the wild-type residue. Together, these results provide strong evidence for strain-specific epistasis at site 83, where the fitness consequences of individual missense substitutions depend on the Vif sequence background. We observed differential effects for some, but not all, Q83 variants between the A3G and A3H conditions. Our findings suggest that these differences may reflect either substrate-specific interactions or context-dependent effects on Vif stability or expression. These findings highlight the strain-dependent nature of adaptive potential at the Vif–A3G interface and underscore how evolutionary trajectories are shaped by genetic context and prior evolutionary history.

## Discussion

Our study provides a comprehensive, high-resolution map of the evolutionary constraints and adaptive potential of HIV-1 Vif, the primary antagonist of the host restriction factor APOBEC3G (A3G) and other APOBEC3 proteins. By applying deep mutational scanning (DMS) to *vif* from two divergent HIV-1 strains, we uncovered both mechanistic insights into Vif function and broader principles governing viral antagonism of host innate immunity. Importantly, we found a strong inverse correlation between variant enrichment scores and A3G-mediated mutation rates, reinforcing DMS-derived enrichment as a highly reliable quantitative proxy for viral fitness in the presence of A3G and enabling the identification of previously uncharacterized, functionally competent Vif variants. More broadly, our approach demonstrates that DMS can be applied to map mutational landscapes of viral accessory proteins even in the presence of host enzymes such as A3G, which introduce mutations independent of the experimental design. Within these landscapes, we observe both pervasive purifying selection and unexpected pockets of mutational tolerance, revealing a balance between functional constraint and evolutionary flexibility in Vif.

In our DMS experiments, many Vif residues exhibited signs of purifying selection. Substitutions at conserved motifs, such as those required for CBFβ binding or within well-characterized functional domains, most severely compromised activity. Yet we also observed pockets of unexpected tolerance, even at residues embedded within A3G, RNA, and CBFβ binding interfaces. These sites highlight Vif’s ability to accommodate alternative sequence states while retaining activity, a hallmark of its evolutionary plasticity.

Crucially, by comparing two Vif proteins shaped by distinct evolutionary contexts and selective pressures, we identified multiple positions that differ in mutational tolerance at binding interfaces with CBFβ, A3G, and RNA, potentially reflecting distinct host pressures that shaped the evolution of LAI and 1203. LAI Vif efficiently antagonizes A3G but not A3H, whereas 1203 Vif can counteract both (*58*). Accordingly, LAI Vif appears optimized primarily for A3G antagonism, whereas 1203 Vif must accommodate dual selective pressure from A3G and A3H. Such expanded functionality may impose additional constraints on Vif, limiting its optimization for any single target and helping explain why mutations tolerated in LAI can be deleterious in 1203, or vice versa. Together, these findings support a model in which host genetic variation shapes strain-specific Vif fitness landscapes and drives divergent evolutionary trajectories. More broadly, Vif proteins across HIV-1 strains can be subject to additional selective pressures beyond A3 proteins, including interactions with host factors such as PPP2R5 (*32–36*). Consistent with prior work showing that PPP2R5 degradation is not uniformly conserved across HIV-1 strains (*32, 36*), neither LAI nor 1203 Vif has been reported to efficiently engage this pathway. Thus, unlike A3H antagonism, differences in PPP2R5 targeting are unlikely to explain the strain-specific effects observed here. However, additional host protein interactions beyond A3G might also influence the enrichment/depletion scores in our assays and may explain selection at sites not known to be involved in the Vif–A3G interaction.

We identified multiple positions where the same substitution produced divergent effects in LAI versus 1203 (Fig. 5). This strain-specific epistasis highlights the dynamic nature of the Vif–A3G arms race, in which mutational effects depend strongly on genetic background. Such differences would not be apparent from analysis of a single strain alone, underscoring the value of comparative DMS approaches for dissecting host–virus conflict. Position 83 provides a vivid example of such dynamics. The ancient Y83H substitution enabled cross-species transmission from simian hosts to humans (*22*). Thus, it is not surprising that H83 is the most common residue found in many non-clade B HIV-1 lineages. We also show that both H83 and Q83 are equally fit in Vif from the clade A HIV-1_Q23-17_ strain (Fig. 7B). However, Q83H is significantly fitness impaired in both clade B viruses, LAI and 1203 (Fig. 6C-E), consistent with its low frequency among clade B isolates (Fig. 7A). These findings provide direct evidence for temporal epistasis, in which residues that were once adaptive become deleterious in a new genetic context. Conversely, Q83C and Q83S enhanced A3G antagonism in LAI but were deleterious or neutral in 1203 (Fig. 6C-E), demonstrating strong strain-specific epistasis even among closely related clade B viruses.

Although our DMS data revealed many variants with positive enrichment scores, most are absent from circulating HIV-1 strains. While some of these may reflect context-specific gains in fitness, others could arise from experimental noise, sampling variance, or artifacts of pooled competition. Accordingly, enriched variants require careful interpretation and independent validation. One notable exception is Q83C, which emerged in our dataset as highly enriched and was previously shown to enhance Vif-mediated antagonism of an SIV from African Green Monkeys A3G that was adapted *in vivo* (*79*). Importantly, Q83C in the context of our HIV-1 experiments is an adaptive mutation with increased A3G antagonism (Fig 6 and 7). Consistent with this finding, we find that H83C also confers a fitness advantage in a clade A Vif background (HIV-1_Q23-17_ strain), demonstrating that this effect is not restricted to laboratory-adapted strains and can extend to more natural viral contexts. It remains unclear whether this activity stems from reinforcement of known contacts at the RNA/A3G binding interfaces or from the emergence of novel interactions, and whether constraints have prevented the emergence of this amino acid change in circulating HIV-1 strains. In contrast to Q83C, enriched variants at other positions, such as site 42, exclude A3G from virion incorporation at levels comparable to wild-type Vif *in vitro* (Fig. 4C) but are not sampled in nature (*23*), suggesting additional constraints not captured by our assays. Together, these findings emphasize that while depleted variants reliably indicate functional disruption, enriched variants should be interpreted cautiously in the absence of orthogonal validation (*82*).

To place these findings in a broader evolutionary context, we compared site-level median enrichment scores from our DMS analysis of A3G antagonism with residue conservation across HIV-1 Group M, including circulating recombinant forms, using the curated Vif subtype alignment from the Los Alamos National Laboratory HIV Sequence Database (Supplementary Data 5). We observed very little correlation between mutational tolerance in our assay and evolutionary conservation, in both LAI and 1203 (Supplementary Fig. 5). This disconnect extended across most residues in both datasets, suggesting that sites tolerant of mutation *in vitro* are often highly conserved *in vivo*. One interpretation is that natural HIV-1 sequences are not at evolutionary equilibrium; strong phylogenetic relatedness can limit observed variability even at mutationally tolerant sites. Indeed, prior DMS studies of HIV Env have similarly reported only modest correlations between experimentally determined amino acid preferences and natural sequence frequencies (*44*). Thus, even when enrichment scores provide a high-resolution, experimentally tractable view of A3G-specific functional constraints, they do not fully explain natural sequence conservation *in vivo*. This discrepancy likely reflects the integration of multiple selective pressures in natural populations, including historical contingency, structural dependencies, evasion of multiple A3 proteins, interactions with cellular cofactors, and constraints imposed by overlapping open-reading-frames.

In the DMS LAI *vif* dataset, sites 13 and 14 exhibited higher-than-average median log enrichment scores (Fig. 3A; Fig. 4A), and in the DMS 1203 *vif* dataset, sites 12,15, and 16 exhibited higher-than-average median log enrichment scores (Fig. 5B; Supplementary Fig. 4), suggesting that several substitutions were tolerated and may even moderately enhance A3G antagonism. This is particularly interesting because, like site 83, position 15 falls within the well-characterized “arms-race” interface between Vif and A3G (*23*). Notably, these residues are almost universally conserved across HIV-1 Group M (Supplementary Data 4). Our results suggest that this conservation is driven primarily by constraints in *pol* rather than intrinsic functional requirements within *vif*. A similar trend appears at position 19, which overlaps with the *pol* stop codon in natural HIV-1 isolates but, like sites 13 and 14, shows higher-than-average median log enrichment in the DMS LAI *vif* library (Fig. 4B). Depending on whether *pol* terminates with TAG, TAA, or TGA, the overlapping *vif* codon could encode AGX (Serine/Arginine), AAX (Asparagine/Lysine), or GAX (Aspartate/Glutamate). However, because TGA is never used as the *pol* stop codon, site 19 in Vif is almost always restricted to lysine (K, 7.5%), asparagine (N, 17.6%), arginine (R, 73.6%), or serine (S, 1.2%) (Supplementary Data 4). In contrast, our DMS data revealed high enrichment scores for several additional substitutions at site 19, suggesting that mutational tolerance at this site increases when the overlap with *pol* is removed. Notably, while aspartate is moderately enriched in the DMS LAI *vif* library (average log enrichment score of 1.72), both aspartate and glutamate are strongly depleted in the DMS 1203 *vif* library (average log enrichment scores of –2.91 and –2.02, respectively; Fig. 5B), consistent with prior evidence that site 19 contributes to the dual A3G–RNA interface (*23*). Together, these observations suggest that selection against acidic residues at Vif site 19 may have simultaneously constrained *pol* from evolving a TGA (opal) stop codon, thereby imposing reciprocal evolutionary constraints on both overlapping reading frames. In contrast, our DMS assay isolates the contribution of A3G antagonism under a controlled genetic background and relieves mutational constraints imposed by the overlapping reading frame shared between *vif* and the C-terminal tail of IN, which is critical for viral integration *in vitro* and *in vivo* (*83, 84*) and for stabilizing the intasome complex (*85*).

Collectively, our findings establish a validated DMS assay for HIV-1 Vif, enabling high-throughput, quantitative analysis of variant fitness in the presence of A3G. By combining pooled functional selections with targeted validation assays, we reveal the delicate balance Vif must strike to maintain effective A3G antagonism while accommodating competing pressures from other host factors such as A3H and from viral genome architecture. Our approach highlights both well-established constrained motifs and previously uncharacterized, functionally competent variants, including Q83C. It also reveals mutational tolerance at positions that are highly conserved, offering insight into evolutionary plasticity not readily accessible through sequence analysis alone. More broadly, this work illustrates how DMS can disentangle individual selective pressures from the complex evolutionary landscape of viral accessory proteins, providing a general framework for identifying functional constraints and evolutionary trade-offs in host–virus interactions.

## Materials and Methods

### Cell culture and transfections

SUPT1 cells stably expressing A3G at physiological levels were a gift from Reuben Harris and Judd Hultquist, as previously described (*3*). These cells were maintained in RPMI-1640 medium (Gibco, #11875093) supplemented with 10% fetal bovine serum (FBS; GE Healthcare, #SH30910.03), 1% penicillin–streptomycin (Gibco, #15140122), and 1% HEPES at 37°C with 5% CO₂. Unmodified SUPT1 cells (ATCC CRL-1942) used in validation assays were cultured in RPMI-1640 supplemented with 10% FBS, 1% Antibiotic–Antimycotic (Gibco, #15240062), 1% sodium pyruvate (Gibco, #11360-070), 1% glucose (Gibco, #A24940-01), and 1% GlutaMAX (Gibco, #35050-061). HEK293T cells (ATCC CRL-3216) were maintained in DMEM (Gibco, #11965092) supplemented with 10% FBS and 1% penicillin–streptomycin. HEK293T-c17 cells (ATCC CRL-11268) were maintained in DMEM with 10% FBS and 1% Antibiotic–Antimycotic.

### Plasmids

The plasmid encoding 1203 Vif was described previously (*58*). Site-directed point mutations for infectivity experiments were introduced using PrimeSTAR HS DNA Polymerase (Takara, #R010B) or synthesized as gene fragments (Twist Bioscience, San Francisco, CA). Mutated fragments were inserted into the HIV-1 LAI-based molecular clone pLAIΔVifΔEnvLuc2 using MluI and XbaI restriction sites, as previously described (*22, 58*). For packaging assays, position 42 variants were cloned into pHIV packaging vectors and co-transfected with psPAX2 and pMD2.G plasmids (gifts from Didier Trono; Addgene plasmids #12260 and #12259, respectively). The codon-mutant DMS libraries for LAI and 1203 Vif were cloned into a replication-competent HIV-1 proviral backbone in which the endogenous vif start codon overlapping with pol was ablated, as described previously (*16*). Libraries were digested with MluI and XbaI and ligated into this backbone to generate DMS LAI Vif HIV-1 and DMS 1203 Vif HIV-1 constructs. All plasmid sequences were confirmed by Sanger sequencing (Fred Hutch Genomics and Bioinformatics Shared Resource) or full plasmid sequencing (Plasmidsaurus).

### Construction of DMS *vif* libraries

Deep mutational scanning (DMS) libraries were designed to encode all possible single amino acid substitutions across residues 12–115 of the *vif* genes from HIV-1 LAI and 1203 strains. For each codon, degenerate sequences were selected to encode the desired substitutions while minimizing the introduction of A3G-preferred deamination motifs (e.g., 5′-CC) when alternative codon choices were available. Oligonucleotide pools were synthesized by Twist Bioscience (San Francisco, CA, USA). Variant codons for each position were synthesized as gene fragments flanked by MluI and XbaI restriction sites and delivered in a 96-well plate format, with each well corresponding to a specific residue. For each site, codons encoding the desired amino acid substitutions were pooled in equal proportions. The variant pools for all targeted positions were then combined in equal amounts to generate the complete DMS *vif* libraries for LAI and 1203. These libraries were subsequently cloned into the replication-competent HIV-1 backbone described above.

### Screening DMS *vif* viruses

For each of three biological replicates, 3.1 × 10⁷ SupT1-A3G cells were infected at an MOI of 0.01 by spinoculation (1100 × g, 30 min) in the presence of 20 μg/mL DEAE-dextran. The media was replaced immediately and again 24 hours post-infection. At 72 hours, the supernatant was harvested, supplemented with 20 μg/mL DEAE-dextran, and used to infect 1.5 × 10⁷ fresh SupT1-A3G cells by spinoculation. Following a second 72-hour infection, the post-selection supernatant was collected. For sequencing, supernatants were filtered (0.2 μm) and concentrated using a SW28 rotor (1 h, 4 °C). Viral pellets were resuspended in PBS, and viral RNA was extracted using the Quick-RNA Viral Kit (Zymo Research, #R1035).

### Sequencing analysis of viral supernatants

Viral RNA was reverse transcribed with SuperScript IV (Thermo Fisher #18090010), and *vif* cDNA was amplified by RT-PCR using primers annealing outside the mutagenized region. Amplicons were processed for library preparation as described in (*45*). Multiplexed libraries were pooled and sequenced on an Illumina NextSeq P2 platform using 250 paired-end reads (Fred Hutch Genomics and Bioinformatics Shared Resource).

### Analysis of DMS results

#### Processing of merged reads and consensus generation

Merged paired-end sequencing reads were processed using custom Python scripts to generate high-confidence consensus sequences for each unique molecular identifier (UMI). Reads were parsed from FASTQ files and filtered to retain only those corresponding to the expected insert length, excluding truncated or over-extended fragments from downstream analysis. For each UMI, nucleotide identities at each position were aggregated across all supporting reads. To generate the consensus call at each base, the nucleotide with the highest associated Phred quality score was selected. This quality-weighted approach prioritizes the most confidently called bases and reduces the influence of low-frequency sequencing errors. The resulting consensus set represented high-confidence full-length coding sequences suitable for codon-level analysis.

#### Generation of codon count tables

High-confidence consensus sequences were translated into codon triplets and mapped to amino acid positions 12–115 of the LAI or 1203 reference Vif sequence. Each codon observed at a given position was tallied to produce codon count tables, where each row corresponded to a codon position in the reference sequence and each column indicated the count of a specific codon variant. The resulting codon count matrices served as the input for subsequent steps of variant identification and enrichment analysis.

#### Site-level enrichment analysis, normalization, and statistical testing

To quantify the selective effects of individual mutations in Vif, codon-level variant counts from pre- and post-selection libraries were converted into relative frequencies. A pseudocount of 0.5 was added to all codon counts prior to frequency normalization to stabilize log ratios for low-count variants and avoid division by zero. For each variant, the change in frequency after selection was measured by comparing its post-selection abundance to its pre-selection abundance. Pre-selection replicates were pooled to generate a single, high-depth baseline library capturing the full diversity of input variants while minimizing sampling noise. During pooling, variants that did not exceed a per-sample background threshold —estimated from non-mutagenized sites (sites 1-11 and 116-122)— in a minimum of two replicates were zeroed out, ensuring that the merged baseline reflects only robustly observed variants. This shared pre-selection reference was used across all post-selection replicates within each strain, ensuring that enrichment calculations reflect biological effects of selection rather than fluctuations in the input distribution. Using a common baseline also improves the statistical power to detect enrichment in low-frequency variants. Because sequencing depth and wild-type representation can vary across codons due to library biases or amplification noise, enrichment ratios were normalized relative to the wild-type codon at each site. This site-specific normalization accounts for local differences in coverage and corrects for site-specific distortions in baseline representation. In effect, each variant’s enrichment score reflects its change relative to the wild-type codon’s behavior under selection. Normalization was only performed at sites where the wild-type codon was sufficiently represented in both pre- and post-selection libraries.

For each strain, enrichment ratios were calculated independently for all biological replicates, using the same pooled pre-selection baseline. At each codon position, variant-level enrichments were aggregated by computing the median enrichment across all missense substitutions, yielding a site-level enrichment score. This approach reduces noise from individual variants and captures the overall mutational tolerance or constraint at each residue.

To identify sites under selection within each strain, all variant-level enrichment values at each site, pooled across replicates, were statistically compared to the global background of enrichment scores using Wilcoxon signed-rank tests. Two complementary tests were performed: (i) a two-sided test to detect sites with any deviation (enrichment or depletion) from the global median, and (ii) one-sided tests to identify sites with consistent enrichment (mutational tolerance) or depletion (functional constraint). False discovery rate (FDR) correction was applied using the Benjamini–Hochberg procedure (FDR ≤ 0.10) to identify statistically significant positions.

#### Identification and quantification of A3G-mutated reads

To identify and quantify A3G-mediated hypermutation in the DMS libraries, consensus FASTQ files were parsed and compared against the reference *vif* sequence in Python. Each read was aligned codon-by-codon to the reference open reading frame, and all single-nucleotide mismatches were recorded along with their position, codon context, and base substitution identity. Substitutions consistent with A3G activity were annotated.

### Functional validation of DMS *vif* variants

Single-cycle infectivity assays were performed as described (*58*). HEK293T-c17 cells were seeded at 1.5 × 10^5^ cells/ml in 6-well plates and transfected with 400 ng pLAIΔVifΔEnvLuc2 proviral vector, 100 ng L-VSV-G, 3XFlag-tagged A3G, HA-tagged A3H Hap II, or pcDNA4/TO empty vector, and Transit-LT1 (Mirus #MIR2300) at 3.5 μL reagent/μg DNA. Media was replaced after 24 hours, and virus was harvested 72 hours post-transfection, then filtered and normalized by RT-qPCR-based titering (*86, 87*). Virus inputs were normalized by mock supernatant addition to equalize volumes. Unmodified SupT1 cells were seeded at 1.5 × 10^5^ cells/well in 96-well plates in medium with 20 μg/ml DEAE-dextran. At 72 hours post-infection, cells were lysed in Bright-Glo reagent (Promega #E2610), and luciferase activity was measured on a LUMIstar Omega luminometer (BMG Labtech). Each condition was measured in duplicate technical replicates. Fold difference in infectivity was calculated by dividing the Raw RLUs by the average RLUs for the relevant wild-type virus (wild-type LAI, wild-type 1203, or wild-type Q23-17). These values were then further normalized by dividing by the mean fold difference in infectivity for *vif* variants tested without any A3 plasmids (Separate experiment, Fig. 6D).

### Packaging assay validation of H42 variants

HEK293T-c17 cells were seeded at 1.5 × 10⁵ cells/mL in 6-well plates and transfected with 500 ng psPAX2, 100 ng 3×Flag-tagged A3G in pcDNA4/TO, 500 ng pHIV lentivector expressing Vif variants or no Vif, and TransIT-LT1 (Mirus #MIR2300) at 3.5 uL reagent/μg DNA. Media was replaced after 24 hours, and virus was harvested 72 hours post-transfection, then filtered and concentrated by centrifugation (1 h, 4 °C, maximum speed). Viral pellets were resuspended in PBS, normalized by HIV-1 p24 ELISA (R&D #DY7360-05), and prepared for SDS-PAGE using NuPAGE loading dye (Invitrogen #NP007). Proteins were resolved by SDS-PAGE, transferred, and immunoblotted with anti-Flag (Sigma #F1804) or anti-p24 (HIV-ARP #3537), followed by anti-mouse IgG secondary (R&D #HAF007), all at 1:5000. Blots were imaged on a ChemiDoc MP (Bio-Rad) and quantified with ImageJ. A3G incorporation was calculated as the anti-Flag/anti-p24 ratio, normalized to the ΔVif control.

### Analysis of primary isolate Vif sequences

Primary isolate sequences were obtained from curated alignments provided by the Los Alamos National Laboratory HIV sequence database. Group M subtype reference alignments including circulating recombinant forms (CRFs) were used for global analyses. For Clade B–specific analyses, sequences were downloaded using the “one sequence per patient” option in the LANL database interface. Percent frequencies were calculated from alignments of these sequences (Supplementary data 4/5)

## Acknowledgments

We thank John Gross, Nicholas Chesarino, and Hugh Haddox for feedback on the manuscript and Julie Overbaugh, as well as other members of the Malik and Emerman labs, for valuable discussions and feedback. We also thank Hugh Haddox for input on initial DMS analysis, Jeannette Tenthorey for input on data analysis and visualization, and the Fred Hutch Genomics Core for sequencing.

## Author contributions

Conceptualization: CAL, HSM, ME

Methodology: CAL, ME, HSM

Investigation: CAL, ML

Visualization: CAL, ML

Supervision: HSM, ME

Writing –original draft: CAL

Writing –review & editing: CAL, HSM, ME, ML

## Data and materials availability

All data are available in the main text or the supplementary materials.

Source code for all analyses and figure generation is publicly available at GitHub (https://github.com/carolangley/DMS-Vif-Analysis).

Raw sequencing data have been deposited in the NCBI Sequence Read Archive (SRA) under submission number SUB15734269 and will be released upon publication.

---

**Fig. S1.**
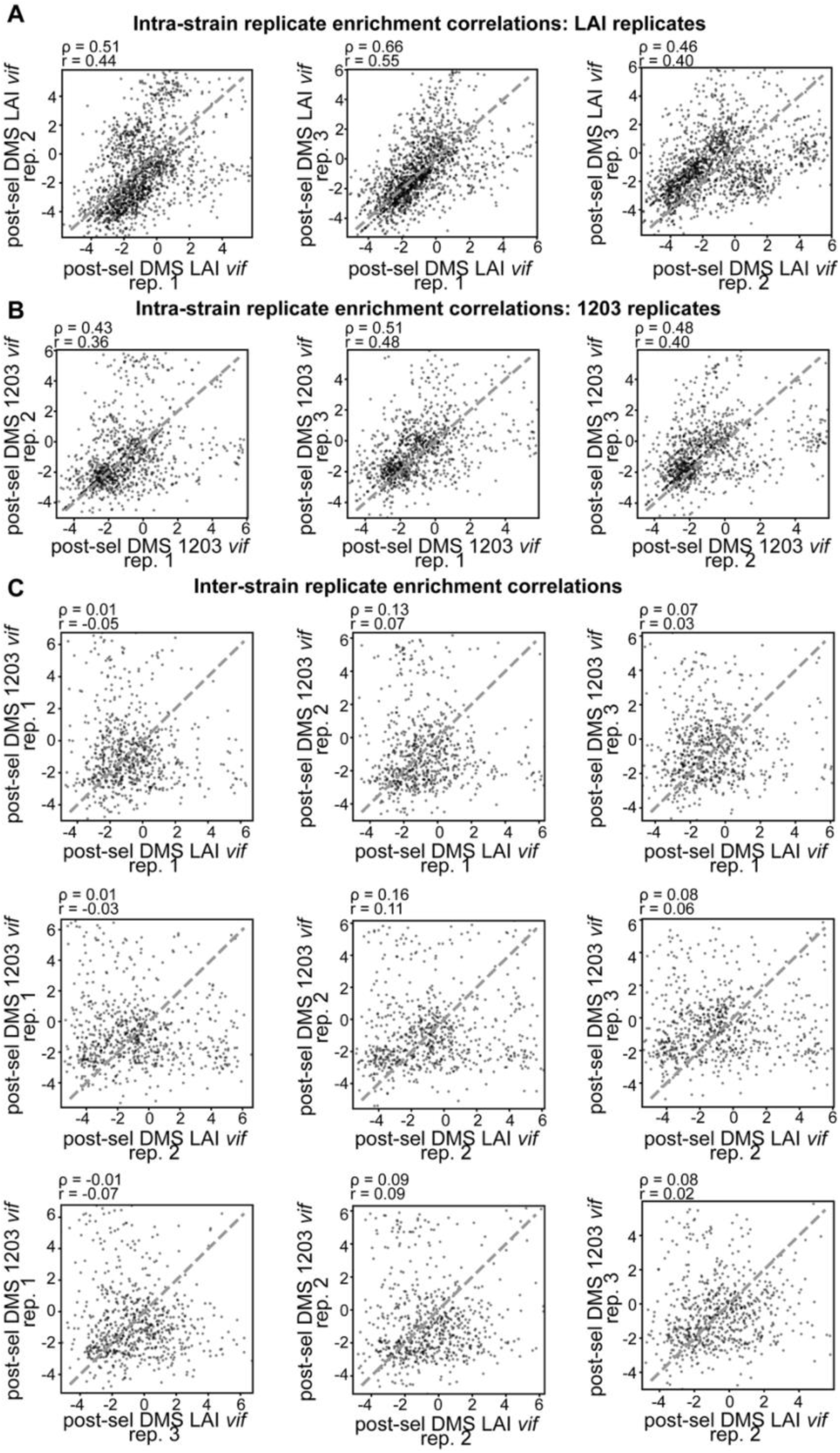
Pairwise Spearman correlations of DMS enrichment scores across replicates and strains. Correlation plots comparing enrichment scores between biological replicates within the same HIV-1 Vif strain (intra-strain) and across different strains (inter-strain). Each point represents a single Vif variant’s enrichment score the DMS assay.

**Fig. S2.**
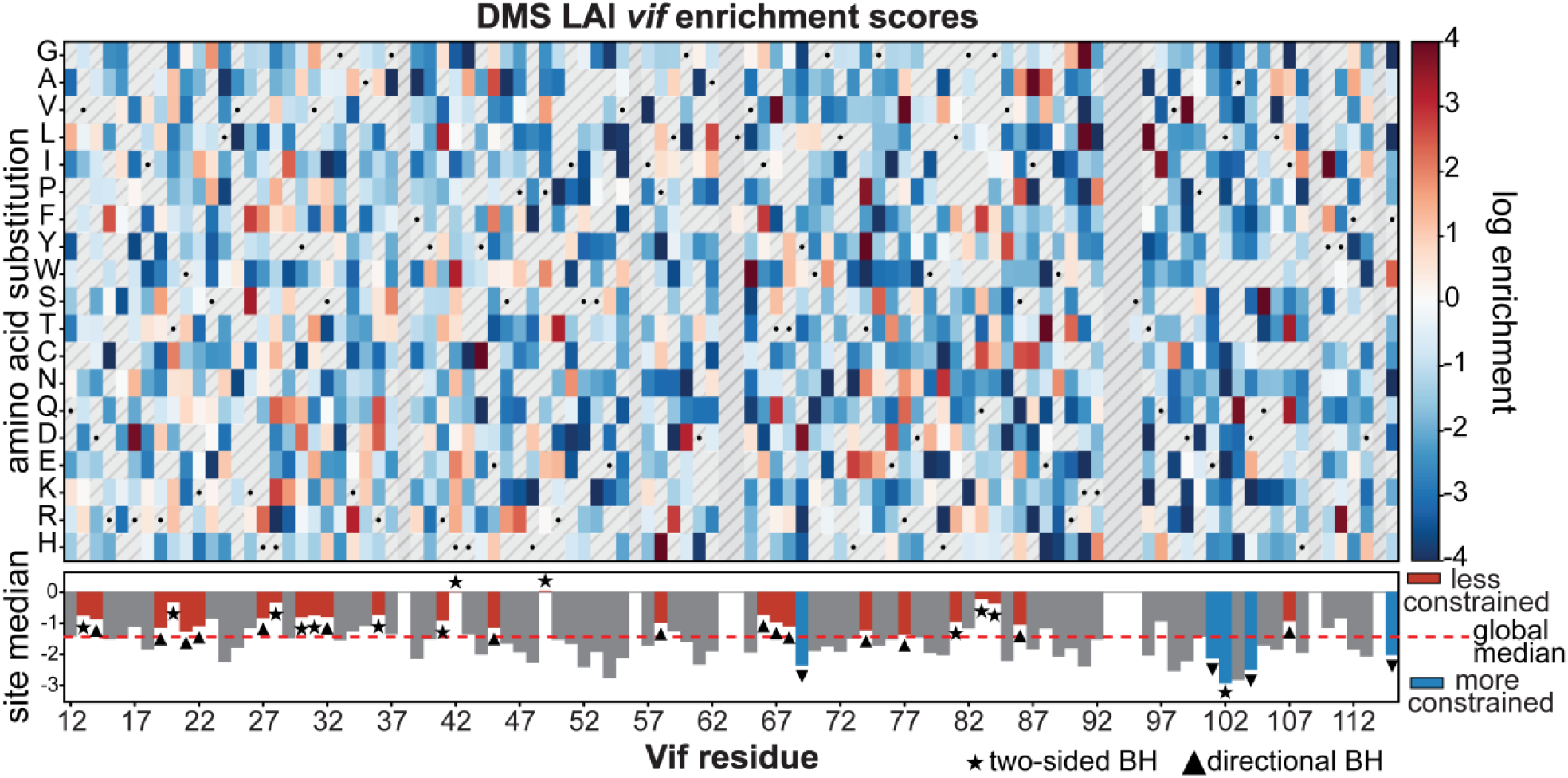
Heatmap showing the log enrichment ratios for all single missense substitutions across residues 12–115 of LAI Vif following two rounds of selection in SupT1 cells expressing A3G. Top: Enrichment values reflect relative variant fitness; positive values (red) indicate retention or enhancement of A3G antagonism, whereas negative values (blue) indicate depletion after selection. Sites with insufficient variant coverage (<5 missense variants) are colored gray and hatched and excluded from statistical analyses. Additionally, sites 38, 56, 63–64, 93–95, 109, and 114 failed library quality control during synthesis and were excluded from analysis. Individual variants with insufficient representation are also shown in hatched gray. The wild-type amino acid at each site is denoted by a black dot. Bottom: Median log enrichment across all variants at each site. Significance was assessed using Wilcoxon signed-rank tests comparing variant enrichment values at each site to the library-wide median (dashed red line). Sites passing a Benjamini–Hochberg false discovery rate threshold (q ≤ 0.10) in either a two-sided test (star) or a directionally consistent one-sided test (triangle) are highlighted as red bars (less constrained than average) or blue bars (more constrained than average). All other sites are shown as gray bar.

**Fig. S3.**
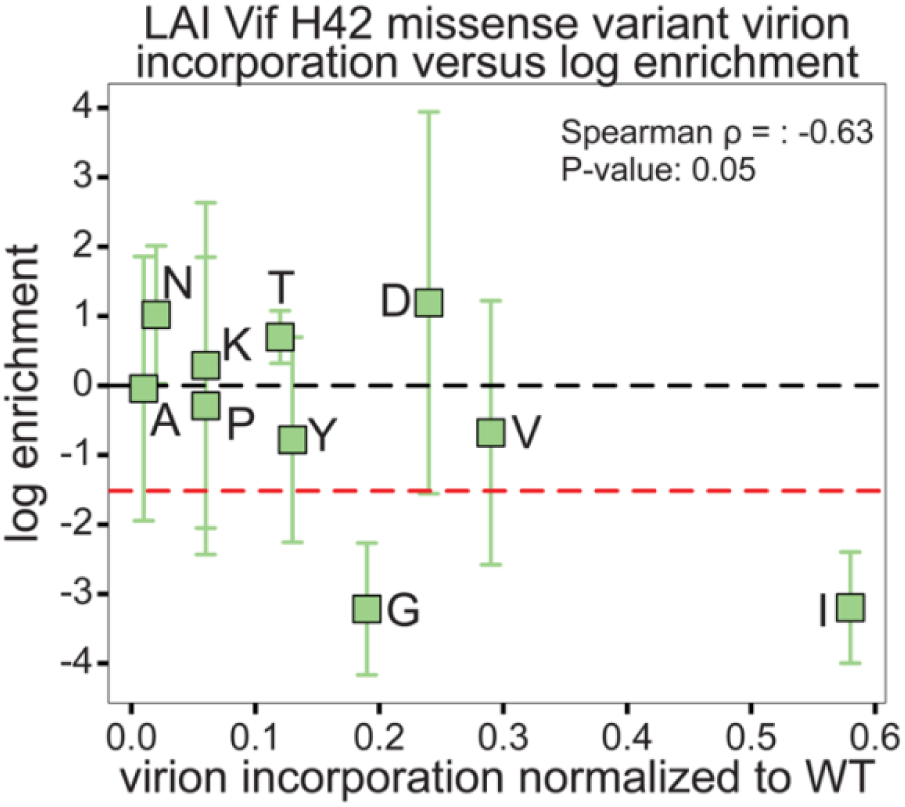
LAI Vif H42 variant enrichment versus virion incorporation. Plot comparing enrichment scores with virion incorporation normalized to the wild-type Vif level for LAI Vif H42 missense variants. Each point represents a single variant and is labeled by the substituted amino acid. Error bars indicate variability in enrichment across replicates. Lower virion incorporation was associated with higher enrichment across variants (Spearman ρ = −0.63, p = 0.05).

**Fig. S4.**
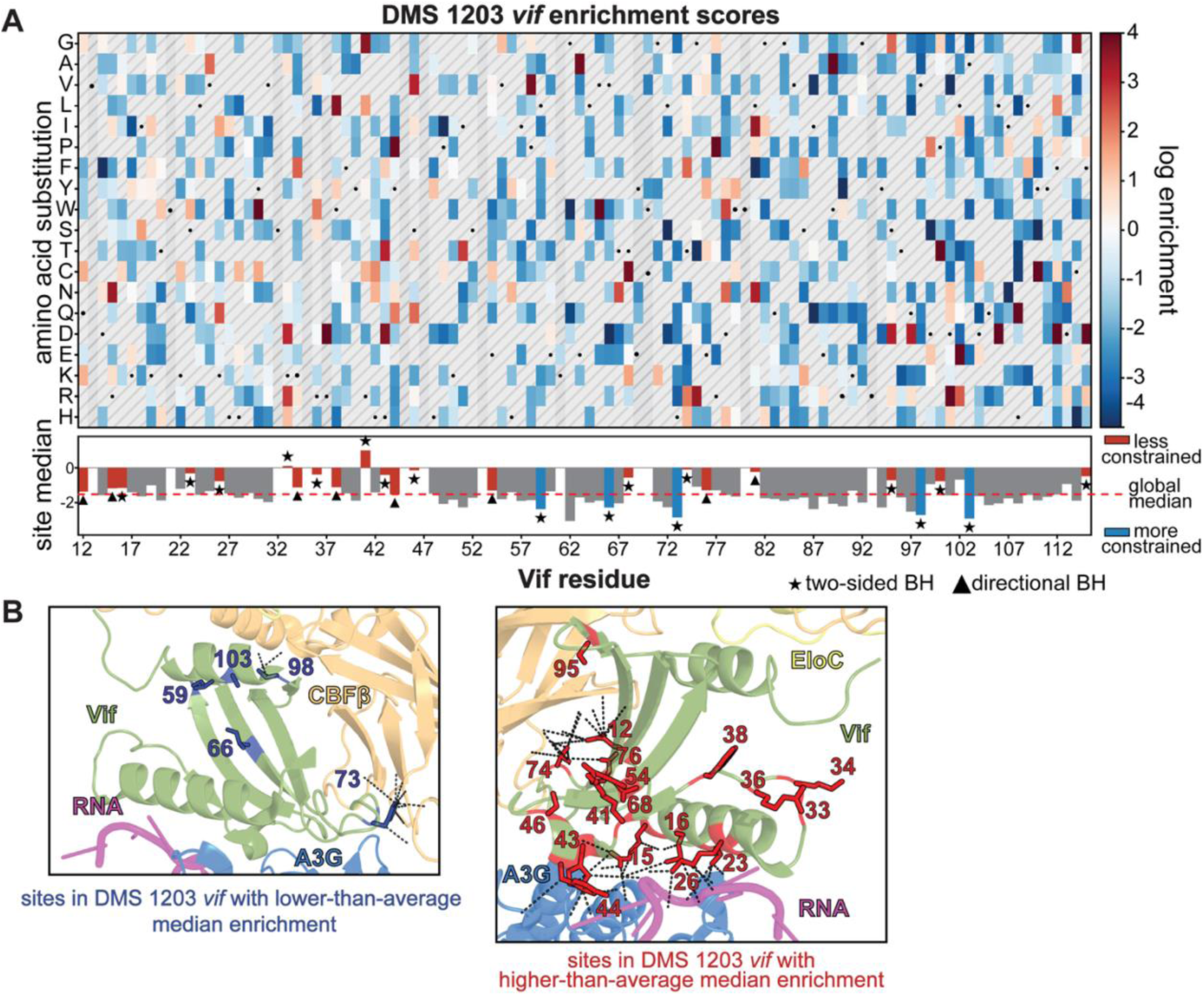
DMS 1203 *vif* dataset heatmap and sites that are significantly more or less constrained than the library-wide average. **A.** Top: Heatmap showing the log enrichment ratios for all single missense substitutions across residues 12–115 of 1203 Vif following two rounds of selection in SupT1 cells expressing A3G. Enrichment values reflect relative variant fitness; positive values (red) indicate retention or enhancement of A3G antagonism, whereas negative values (blue) indicate depletion after selection. Sites with insufficient variant coverage (<5 missense variants) are colored gray and hatched and excluded from statistical analyses. Additionally, sites 21, 35, 45, 47, 53, and 93 failed library quality control during synthesis and were excluded from analysis. Individual variants with insufficient representation are also shown in gray and hatched. The wild-type amino acid at each site is denoted by a black dot. Bottom: Median log enrichment across all variants at each site. Significance was assessed using Wilcoxon signed-rank tests comparing variant enrichment values at each site to the library-wide median (dashed red line). Sites passing a Benjamini–Hochberg false discovery rate threshold (q ≤ 0.10) in either a two-sided test (star) or a directionally consistent one-sided test (triangle) are highlighted as red bars (less constrained than average) or blue bars (more constrained than average). All other sites are shown as gray bars. B. Sites with significantly lower-than-average (left) and higher-than-average (right) median log enrichment scores (from Fig. 5B, bottom panel, blue and red bars) are mapped onto the LAI Vif–A3G–VCBC–RNA cryo-EM structure (PDB: 8CX0). Gray dashed lines indicate known protein–protein and protein–RNA interactions.

**Fig. S5.**
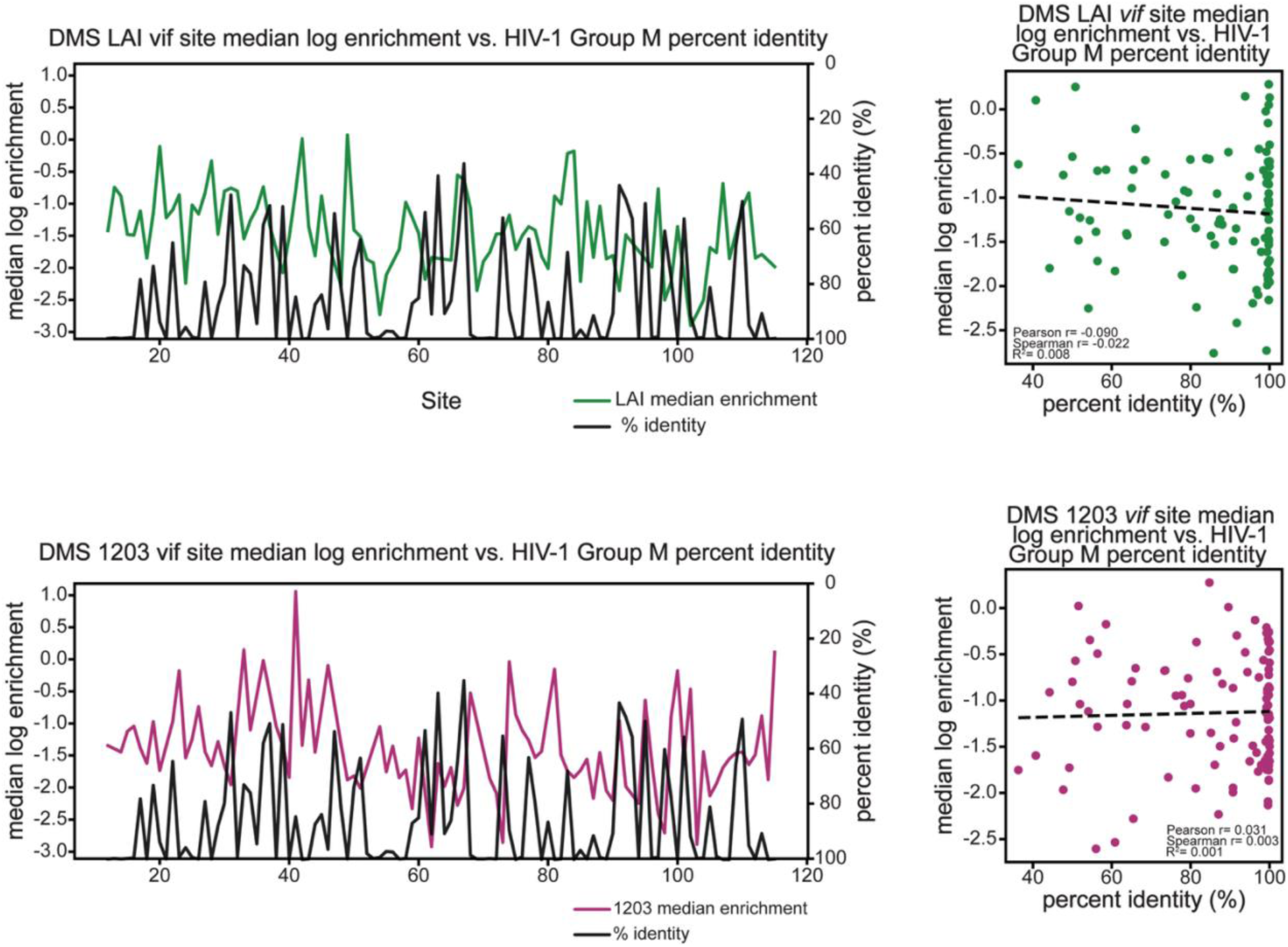
Comparison of functional constraint (via DMS) and evolutionary conservation in HIV-1 Vif. Per-site median log enrichment scores from deep mutational scanning (DMS) of HIV-1 Vif (strain LAI, top row; strain 1203, bottom row) were compared against natural sequence conservation across HIV-1 Group M sequences. Left panels: Median log enrichment values (green for LAI, purple for 1203) are plotted residue by residue across Vif (sites 12–115) alongside percent identity relative to the modal (consnsus) amino acid at each site in a Group M alignment (black line, right y-axis). Right panels: Scatterplots of per-site median enrichment versus consensus percent identity. Each dot corresponds to one Vif residue. Linear regression lines are shown (dashed black), along with correlation statistics (Pearson’s r, Spearman’s ρ). These analyses test whether functional constraint observed in the DMS assays corresponds to evolutionary conservation across natural HIV-1 sequences.

**Data S1. DMS LAI *vif* library QC and variant proportions**

Tabulated summary showing site-level and variant-level QC thresholds, variant codons, the number of sites analyzed, and counts of passed and failed sites.

**Data S2. DMS 1203 *vif* library QC and variant proportions**

Tabulated summary showing site-level and variant-level QC thresholds, variant codons, the number of sites analyzed, and counts of passed and failed sites.

**Data S3. Enrichment ratio metrics for DMS *vif* library variants**

Comprehensive table reporting variant-level enrichment scores from the DMS *vif* experiment. For each mutation, codon-level and amino acid-level substitutions are shown alongside their observed pre- and post-selection counts and frequencies.

**Data S4. Group M Vif Subtype Reference Alignment**

Sequence alignment of representative Vif sequences from Group M viruses

**Data S5. Clade B Vif Subtype Reference Alignment**

Sequence alignment of representative Vif sequences from Clade B viruses

